# Single-cell metabolite detection and genomics reveals uncultivated talented producer

**DOI:** 10.1101/2021.09.11.459929

**Authors:** Masato Kogawa, Rimi Miyaoka, Franziska Hemmerling, Masahiro Ando, Kei Yura, Keigo Ide, Yohei Nishikawa, Masahito Hosokawa, Yuji Ise, Jackson K. B. Cahn, Kentaro Takada, Shigeki Matsunaga, Tetsushi Mori, Jörn Piel, Haruko Takeyama

## Abstract

The production of bioactive metabolites is increasingly recognized as an important function of host-associated bacteria. An example is defensive symbiosis that might account for much of the chemical richness of marine invertebrates including sponges (Porifera), one of the oldest metazoans. However, as most complex microbiomes remain largely uncultivated and lack reference genomes, unequivocally linking metabolic functions to a cellular source is a challenge. Here we report an analysis pipeline of microfluidic encapsulation, Raman microscopy, and integrated digital genomics (MERMAID) for an efficient identification of uncultivated producers. We applied this method to the chemically rich bacteriosponge *Theonella swinhoei*, previously shown to contain ‘Entotheonella’ symbionts providing most of its bioactive substances except for the antifungal aurantosides that lacked biosynthetic gene candidates in the metagenome. Raman-guided single-bacterial analysis and sequencing revealed a cryptic, distinct multiproducer, ‘*Candidatus* Poriflexus aureus’ from a new Chloroflexi lineage. Its exceptionally large genome contains numerous biosynthetic loci and suggested an even higher chemical richness of this sponge than previously appreciated. This study highlights the importance of complementary technologies to uncover microbiome functions, reveals remarkable parallels between distantly related symbionts of the same host, and adds functional support for diverse chemically prolific lineages being present in microbial dark matter.

**Significance Statement:** The production of bioactive metabolites is increasingly recognized as an important function of host-associated bacteria. However, the acquisition of integrated genomic and metabolic data from uncultivated environmental bacteria is still challenging. In this work, we explored the combination of Raman microscopy and single-cell sequencing to localize chemical features to a specific bacterium in an uncultivated microbiome, and we specified the bacteria in the uncultured lineage as a producer of aurantoside, an antifungal natural product, from a chemically and microbially complex sponge. This study offers a new methodology as well as insights into chemical functions of uncultivated life.

## Introduction

Sponges (Porifera) are among the oldest metazoans and thrive in a wide variety of habitats from warm tropical coral reefs to cold deep oceans and freshwater lakes and streams. Their ability to process and filter massive volumes of water during feeding casts a huge impact on biogeochemical cycling^1^ and exposes sponges to a wide range of microorganisms^2^. Many microbes escape lysis by sponges and establish microbiomes that vary from few bacteria to spectacular consortia comprising a major portion of the host biomass. With evidence for a wide range of functional roles that include the acquisition of nutrients, stabilization and reshaping of the sponge skeleton, and disposal of metabolic waste and toxic compounds^3,4^, sponge-microbe associations might contribute to the success of their evolutionarily ancient animal hosts in multiple ways. Characterizing such roles and identifying key microbiota, however, remains a major challenge due to a high complexity of many sponge microbiomes and low success rates of cultivation^5^.

Many sponges contain rich sets of bioactive metabolites with demonstrated or proposed roles in chemical protection and colonization^6^, and as sources of new therapeutics^7^. Recent studies, enabled by metagenomic methods, have uncovered key roles of some sponge symbionts in synthesizing specialized metabolites for their hosts^8–11^. A remarkable example is the chemically rich sponge *Theonella swinhoei* (Fig. 1a), in which a single to few ‘*Candidatus* Entotheonella’ producers within highly diverse microbiomes generate numerous bioactive compounds in their hosts^9,12,13^. ‘Entotheonella’ are members of the new candidate phylum ‘Tectomicrobia’ (Entotheonellaeota)^8^ and form large multicellular filaments that can be enriched by mechanical fractionation^14^. Metagenomic and single-bacterial sequencing of such enriched microbiomes (Fig. 1a) revealed in several *T. swinhoei* chemotypes large sets of biosynthetic gene clusters in ‘Entotheonella’ genomes that were connected to virtually all known sponge polyketides and peptides^8,9^ (Fig. 1b). Besides suggesting important and potentially widespread contributions by sponge microbiota to host defenses, the results demonstrated that uncultivated life harbors lineages with a chemical richness comparable to that of well-known “talented” and industrially important producer taxa, such as filamentous actinomycetes^15^.

**Figure. 1:**
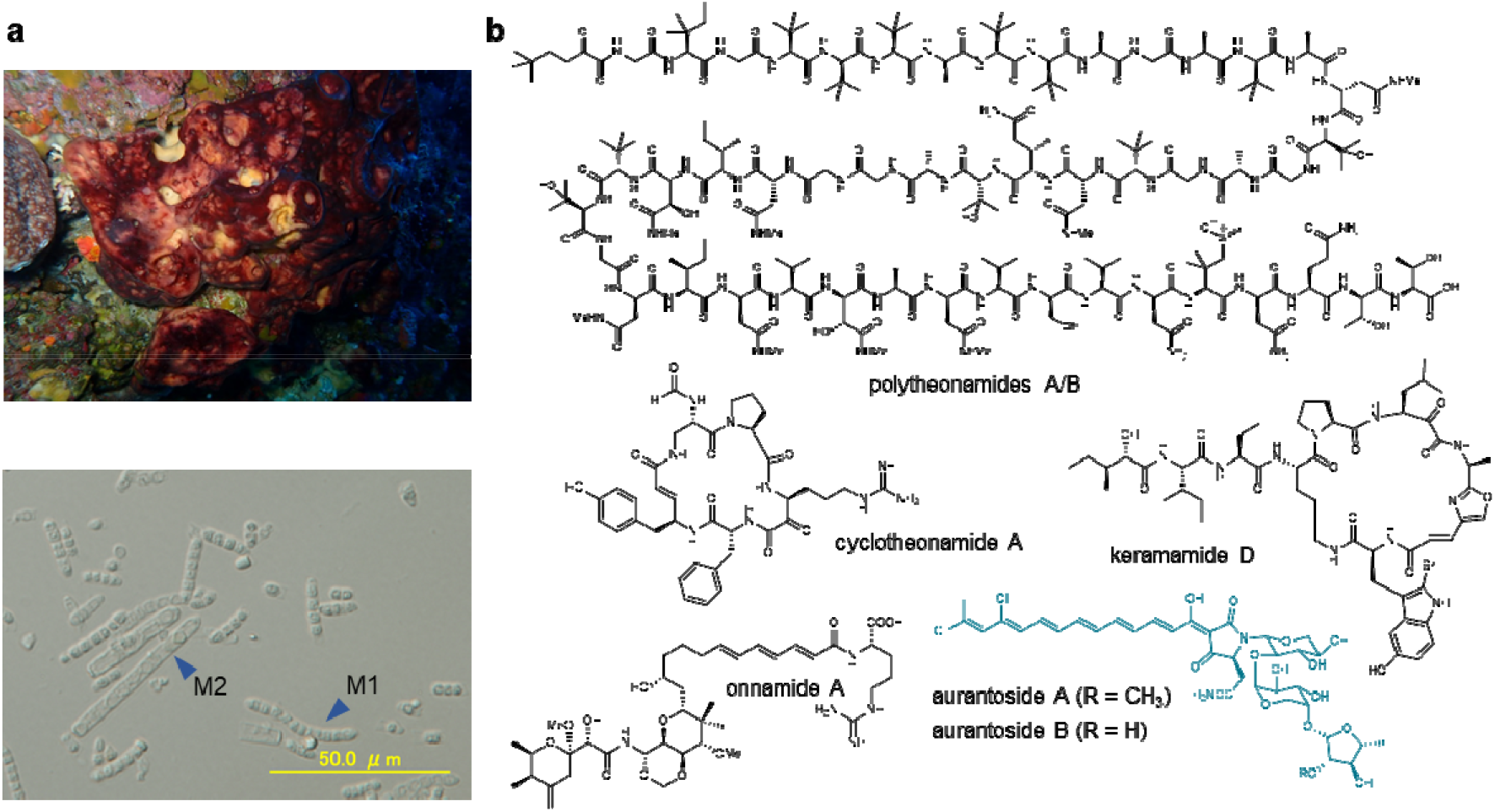
Natural products and filamentous bacterial symbionts of the marine sponge *Theonella swinhoei* TSY. **a)** *T. swinhoei* TSY (top) and a light micrograph of its enriched filamentous symbiont fraction containing two different morphotypes M1 and M2 (bottom). **b)** Structures of secondary metabolites extracted from *T. swinhoei* TSY. Aurantosides are shown in blue.

The identification of metabolic sources in complex microbiomes remains a formidable task. While total metagenomic sequencing can cover much of the sequence space, contigs harboring biosynthetic genes often remain unassigned due to lack of informative taxonomic marker genes or to divergent codon usage that prevent binning. Single-cell genomics, on the other hand, can provide this information but might require the analysis of large sample numbers until a target cell is identified. Here we asked whether an analysis pipeline of **m**icrofluidic **e**ncapsulation, **R**aman **m**icroscopy, **a**nd **i**ntegrated **d**igital genomics (MERMAID) can accelerate the search for key chemical contributors. Tested on a case for which previous methods had revealed neither the organism nor biosynthetic genes, the process rapidly identified a cryptic producer of antifungal metabolites in *T. swinhoei*. The symbiont, a member of a new, deep-rooting *Chloroflexi* lineage that is morphologically similar to ‘Entotheonella’, is highly abundant in the filamentous bacterial fraction but has eluded metagenomic sequencing. On its exceptionally large 14 Mbp genome, numerous biosynthetic gene clusters (BGCs) in addition to the aurantoside locus were identified, suggesting an even higher potential for metabolic richness of this sponge than previously appreciated. Our study highlights the importance of complementary technologies to comprehensively study microbiomes, reveals remarkable parallels between distantly related symbionts of the same host, and adds functional support for diverse chemically prolific lineages present in microbial dark matter.

## Results

### An elusive biosynthetic source of aurantosides in *Theonella swinhoei* Y

Among the diverse characterized bioactive polyketides and peptides known from *Theonella. swinhoei* Y (“Y” refers to the chemotype with yellow interior) (Fig. 1b), the aurantosides represented an exception as the only compounds that could not be assigned to ‘Entotheonella’^8^. Aurantosides comprise a group of yellow-pigmented, structurally related halogenated tetramic acid glycosides with selective antifungal^16^ or cytotoxic^17^ activities. Based on the aurantoside structure, we expected the pathway to involve a polyketide synthase-nonribosomal peptide synthetase (PKS-NRPS) for the polyene-tetramic acid core structure, one or more halogenases, and enzymes for sugar biosynthesis and attachment. In the prior studies on *T. swinhoei* Y, metagenomic binning analyses showed that its mechanically enriched filamentous bacterial fraction contained two ‘Entotheonella’ variants, ‘Entotheonella factor’ and ‘Entotheonella gemina’, with distinct sets of BGCs. These variants appeared to correlate with the presence of two related but discriminable filament morphotypes in the bacterial preparation (Fig. 1a) that had been assumed to represent the ‘Entotheonella’ variants. However, efforts failed to identify matching gene candidates for aurantoside biosynthesis in the BGC-rich ‘Entotheonella’ genomes or the highly fragmented non-’Entotheonella’ metagenomic bins.

When conducting new cell-separation studies with freshly collected *T. swinhoei* Y specimens, we noticed the presence of large yellow bacterial filaments that lost pigmentation after several washing steps with artificial sea water (ASW). In the bleached state, the bacteria appeared identical to the morphotype that we had previously assumed to be one of the two ‘Entotheonella’ variants present in the sponge. Since the ‘Entotheonella’ genomes lacked discernible candidates for an aurantoside BGC but contained genes for carotenoid biosynthesis^18^, the identity of the pigment remained unresolved. This conundrum represented an ideal test case for an integrated Raman microscopy-single bacterial genomics approach.

### The MERMAID single-cell analysis platform

The MERMAID workflow (Fig. 2) starts with single-bacterial encapsulation into microdroplets. For this purpose, a bacterial suspension is introduced into a previously fabricated microfluidic device^19^, and droplets containing single bacteria are generated by dispersion in an oil phase. Then, droplets are isolated into multi-well plates with small contamination risk, and Raman spectra of single bacteria are measured in each well. Finally, bacteria that show Raman bands matching to target metabolites are selected and subjected to single-cell genome amplification in the well. By next-generation sequencing of the amplified genome, selective genome analysis of bacteria containing characteristic metabolites can be achieved.

**Figure. 2:**
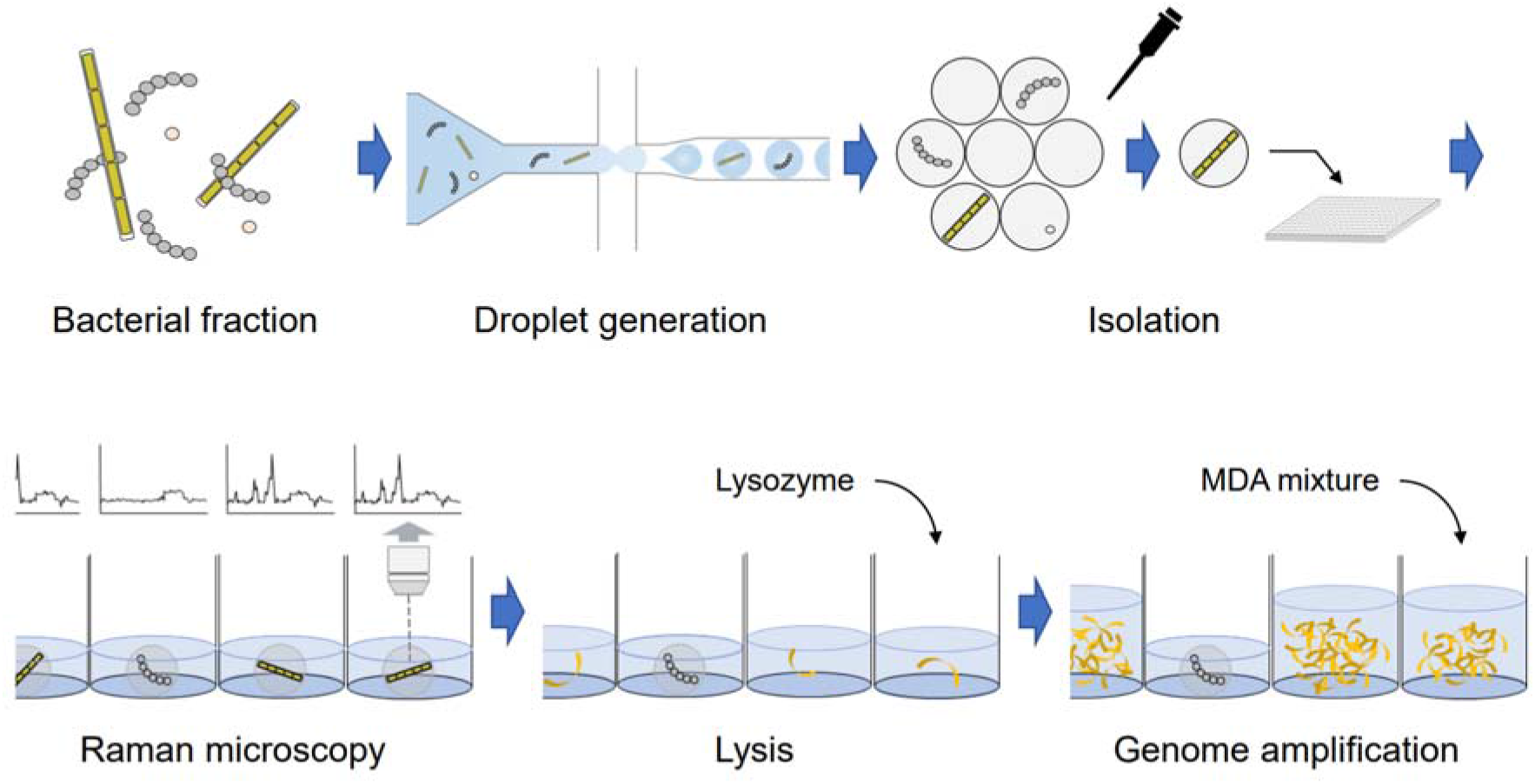
MERMAID workflow. The droplets containing single bacterial filaments are isolated into the well, and Raman spectra of encapsulated bacteria are measured. After droplet disruption by plate shake, cell lysis and genome amplification are implemented in the well.

### Raman microscopy-based identification of an aurantoside producer candidate

To identify the aurantoside source in the microbiome of *T. swinhoei* Y, we initiated this study by acquiring a Raman spectrum from a standard sample of aurantoside A, a chlorinated polyene tetramic acid containing D-xylopyranose, D-arabinopyranose, and C2-methylated 5-deoxyarabinofuranose residues^17^ (Fig. 3a). The spectrum showed characteristic bands at 1,547 cm^−1^ and 1,149 cm^−1^ corresponding to the C=C stretch and the C-C stretch in the hydrophobic polyene moiety.

**Figure. 3:**
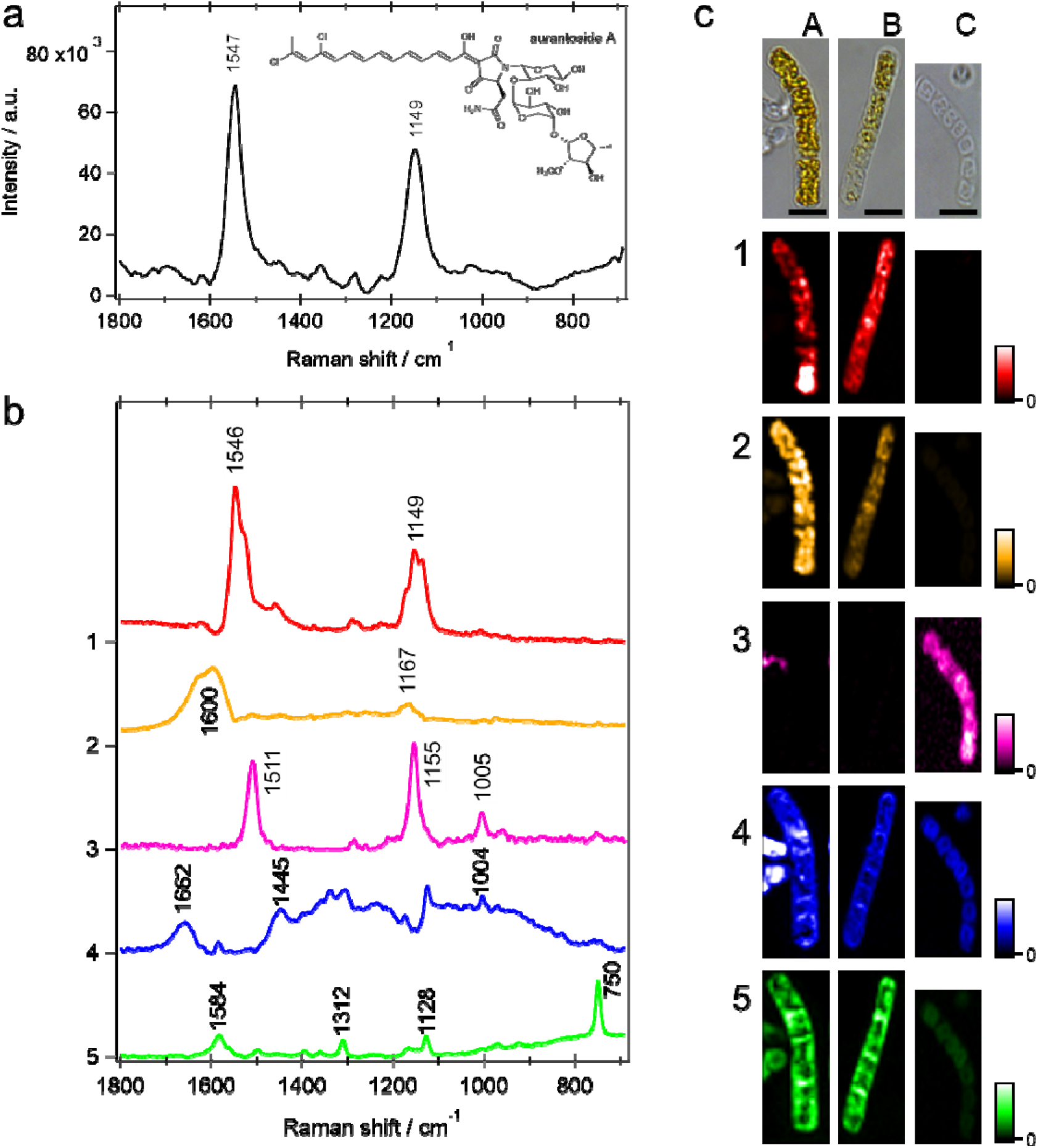
**a)** Raman spectrum of an aurantoside A standard. **b)** MCR-ALS decomposed spectra of bacterial filaments associated with *T. swinhoei* Y. **c)** Phase contrast and MCR-ALS-decomposed images of bacterial filaments (column 1 and 2: ‘P. aureus’ TSY, 3: ‘Entotheonella’). (component 1: aurantoside A, 2: putative aurantoside derivative with shorter polyene length, 3: carotenoid, 4: protein, 5: cytochromes B and C, scale bar: 5 µm)

For single-filament Raman analyses, *T. swinhoei* Y-associated filamentous bacteria were segregated from the sponge tissue by centrifugation, and Raman mapping measurements of single filaments were conducted using the bacterial suspension directly without prior sorting. Subsequently, by applying a multivariate analysis, namely multivariate curve resolution-alternating least squares (MCR-ALS)^20^, to all the spectral data obtained from all the measured filaments, we obtained information on several molecular components. Fig. 3b shows the MCR-ALS-decomposed spectra. Component 1 shows the characteristic bands at 1,546 and 1,149 cm^-1^, which can be assigned to aurantosides. Component 2 shows a similar spectral feature as component 1, but the peaks have shifted to 1,600 and 1,167 cm^-1^. Polyenes show peak shifts of C=C stretch and C-C stretch bands, depending on the length of the conjugated double bonds^21,22^. Therefore, component 2 is assigned to a molecule with shorter polyene length inside the filaments. On the other hand, component 3 is assigned to a carotenoid based on 1511, 1155, and 1005 cm^-1^ bands that correspond to C=C stretch, C-C stretch, and methyl rock, respectively. Among these bands, the band at 1005 cm^-1^ was observed only in component 3, but not in components 1 and 2, consistent with carotenoids carrying methyl groups in the conjugated chain^21,22^ in contrast to aurantosides. Components 4 and 5 are assigned to common proteins and cytochromes B and C, as discussed in our previous study^23^. The MCR-ALS-decomposed Raman images revealed a subset of filamentous bacteria as strong candidates for aurantoside production. As shown in Fig. 3c, aurantosides were only detected in the yellow-colored filaments (columns A and B), but not in a second subset of almost colorless filaments (column C). On the other hand, this colorless phenotype exclusively generated the carotenoid-type spectrum of component 3. Proteins and cytochromes were distributed throughout the cells in all the filaments.

With MCR-ALS-extracted spectra as references, we next tested whether it is possible to establish a microbial characterization pipeline that combines Raman microscopy with single-bacterial sequencing and thus directly links metabolomes to genomes. Raman spectra of droplet-encapsulated single filaments isolated in 384 well plates were individually obtained and analyzed as the linear combination of reference spectra. With an analysis time of a few minutes per well including focusing the cell and 1-second laser irradiation, the method rapidly provided multiple single-bacterial filaments with aurantoside spectral features, thus enabling efficient single-bacterial screening based on metabolite content.

### Single-filament sequencing and genome assembly reveals a cryptic member of the *T. swinhoei* Y microbiome

All aurantoside-positive bacteria identified by Raman microscopy exhibited a uniform morphology consisting of large multicellular filaments comparable in shape to those of ‘Entotheonella’. Characteristic features distinct from ‘Entotheonella’ include a sheath-like structure enclosing each organism and a larger size. To further characterize these bacteria, genomes of 12 filaments with matching Raman spectra were amplified and individually sequenced by Illumina MiSeq, generating a total of 5.25 Gb data at medium quality (>=50% completeness, <10% contamination, Table S1). To construct a high-quality draft genome, the sequence data from the 12 filaments were co-assembled by SPAdes and ccSAG, and the acquired genome was evaluated by QUAST, CheckM, and Prokka^24–28^ (Table S2). The first draft consisted of 3,409 contigs with an overall GC content of 49.7% and was estimated to show 95.6% completeness and 6.2% redundancy based on single-copy marker gene analysis. One rRNA operon and 41 tRNA genes were recovered (Table S3). A BLAST analysis of the 16S rRNA gene sequence retrieved a distant homolog (88.9% identity) from the *Chloroflexi* member *Caldilinea aerophila* (NR_074397) as the closest relative registered in the NCBI 16S RefSeq nucleotide database. When reanalyzing previous metagenomic datasets of the filamentous preparation^8^, we found that the genome of this organism was poorly covered by short contigs despite its abundance and had therefore remained unnoticed. The bacterium was named ‘*Candidatus* Poriflexus aureus’ TSY due to its association with Porifera and its yellow color.

With 14 Mbp, the calculated size of the draft genome exceeded even the large 9-10 Mbp ‘Entotheonella’ genomes and is one of the largest in the Chloroflexi genomes (Table S4), but it is definitely the largest among the recently reported 1,200 sponge symbionts^29^. Since this piece of data appeared highly unusual, we further verified the genome quality by the following investigations (Fig. S1): Individual size estimations conducted from each single-amplified genome (SAG), binning of the contigs based on sequence depths using newly acquired metagenomic shotgun sequencing data, and taxonomic annotation of each contig. Metagenomic sequencing was accomplished with a modified, gentler bacterial lysis protocol after noticing that the *Chloroflexi* filaments rapidly disintegrated during DNA preparation in comparison to ‘Entotheonella’. The size and high quality of the draft genome was confirmed by these analyses, making this one of the largest bacterial genomes reported to date. To further improve the assembly and generate longer contigs for BGC studies, long-read sequencing of SAG mixture was conducted using MinION nanopore technology. By hybrid assembly using long reads and SAG short reads, 195 contigs including one plasmid with a contig N50 of 102,784 bp were acquired (Table S2, Fig. S2). In summary, the data obtained by Raman microscopy and construction of multiple SAGs of ‘P. aureus’ TSY provide evidence for a cryptic filamentous aurantoside producer with a large 14 Mbp draft genome and at least one plasmid.

To test whether ‘P. aureus’ phylotypes are present in other *Theonella* sponges, we investigated a blue specimen of unknown species affiliation containing aurantosides that had been collected around Shimoji island 1,700 km away from Hachijo-jima island (Fig. S3), the collection site of *T. swinhoei* Y. In agreement with our previous data, cell dissociation experiments revealed sheathed filaments that looked identical to those of *T. swinhoei* (Fig. S3). Raman spectra obtained from isolated filaments of the blue sponge were analyzed by MCR-ALS analysis (Fig. S3). In addition to a congener with an aurantoside A-type chromophore, MCR-ALS decomposition suggested the presence of a variant with shortened polyene system based on band shifts to higher frequency. A draft genome of this symbiont was acquired with 68.2% completeness (Table S2). The nucleotide identity of its 16S rRNA gene to that of ‘P. aureus’ TSY was more than 99%. Likewise, the average nucleotide identity of the two draft genomes was calculated as 98.0%, suggesting that both organisms belong to the same candidate species. Because the estimated genome size of the newly identified symbiont was 12.9 Mbp, the large genome was further supported. The symbiont from the blue *Theonella* sp. will be referred to as ‘P. aureus’ BT hereafter.

### Identification and functional analysis of the aurantoside biosynthetic gene cluster

Based on our biosynthetic hypothesis for aurantosides, we searched the ‘P. aureus’ TSY genome for matching gene candidates. This analysis retrieved two small BGC fragments that each contained one halogenase gene, a more extended contig encoding a portion of a trimodular PKS-NRPS, and a contig harboring genes for a putative *S*-adenosylmethionine-(SAM-)dependent methyltransferase and for sugar biosynthesis and attachment. Combinatorial PCR experiments connected these fragments, confirming that they belong to the same locus. Further extension was achieved by MinION sequencing, resulting in a candidate aurantoside (*ats*) BGC with 17 ORFs and of 38 kbp length (Fig. 4a, Table S5).

**Figure. 4:**
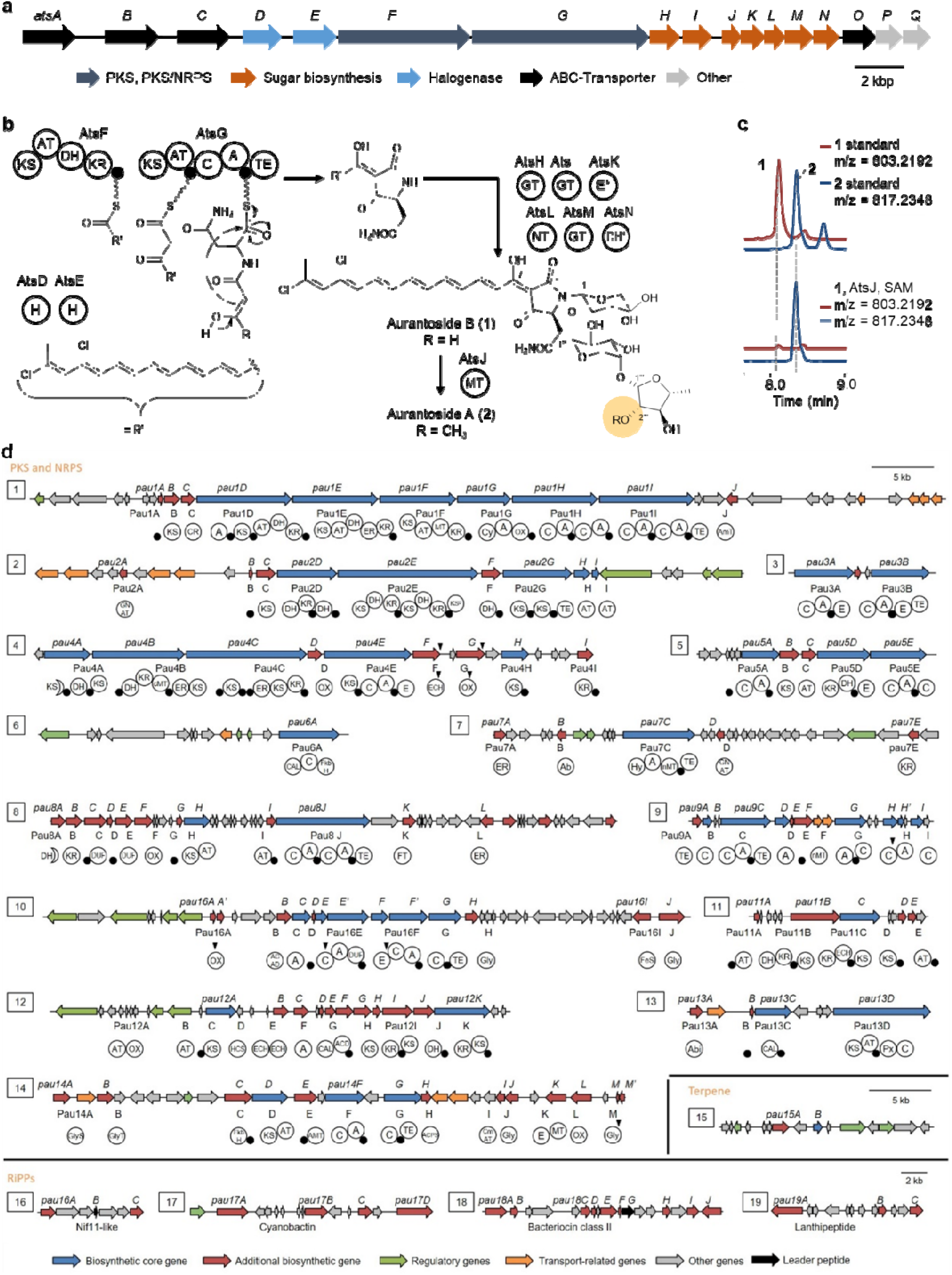
Insights into BGCs encoded in the ‘P. aureus’ genome. **a)** Organization of the aurantoside (*ats*) gene cluster. **b)** Putative biosynthetic pathway for aurantoside A (**2**). The assignments for AtsK and AtsN, marked with an asterisk (*), as sugar epimerases or dehydratases are speculative. The residue modified by AtsJ is highlighted in orange. KS: ketosynthase, AT: acyl transferase, DH: dehydratase, KR: ketoreductase, C: condensation, A: adenylation, TE: thioesterase, H: halogenase, GT: glycosyl transferase, E: epimerase, NT: nucleotidyl transferase, DH: dehydratase, MT: methyl transferase, small black spheres: acyl/peptidyl carrier protein. **c)** Biochemical assays with the *O*-methyltransferase AtsJ. Extracted ion chromatograms are shown for authentic standards of **1** (red) and **2** (blue) (top) and for the conversion of **1** to **2** by AtsJ (bottom). SAM: *S*-adenosyl methionine. For control experiments, see figure S4. **d)** BGCs encoded in the ‘P. aureus’ hybrid draft genome. Triangles indicate stop-codons and frame shifts in the ‘P. aureus’ loci (*pau*) that could not be resolved by resequencing. KS: ketosynthase, CR: crotonyl-CoA reductase, A: adenylation, AT: acyl-transferase, DH: dehydratase, ER: enoyl reductase, KR: ketoreductase, MT: methyl transferase, Cy: cyclase, OX: oxidoreductase, C: condensation, TE: thioesterase, spheres: acyl/peptidyl carrier protein, GNAT: GCN5-related *N*-acetyltransferase, E: epimerase, ECH: enoyl-CoA hydratase, CAL: CoA ligase, Hy: hydrolase, DUF: domain of unknown function, ACD: acyl-CoA dehydrogenase, HCS: holocarboxylase synthase, Px: pyridoxal phosphate-dependent enzyme, AMT: amino transferase. Predicted substrates are noted below the respective A or AT domains.

In order to functionally test the involvement of the *ats* BGC in aurantoside production, we first attempted to perform heterologous expression experiments on the PKS protein AtsF with or without the halogenases AtsD or AtsE. Since none of these trials yielded soluble proteins, we next focused on the methyltransferase homolog AtsJ sharing the highest similarity to the members of the FkbM family (Table S5) that catalyze various *O*-methylations, such as in the polyketide pathways for the immunosuppressant FK506^30^ and the lobophorins^31^. As the only methyltransferase encoded in the BGC, AtsJ represented a strong candidate to convert aurantoside B (**1**) into its *O*-methylated congener aurantoside A (**2**) (Fig. 4b). Initial screens were conducted in cell-free assays with recombinant AtsJ and an authentic standard of **1** isolated from *T. swinhoei* Y. Subsequent assays with purified enzyme contained AtsJ in assay buffer, SAM, and **1**. We observed the conversion of **1** into **2** only in the presence of AtsJ (Fig. 4c), supporting a role of the identified BGC in aurantoside biosynthesis. Controls without SAM, without AtsJ, or with boiled enzyme gave no conversion, while controls without the cofactor Mg^2+^ gave partial conversions, presumably due to the presence of low Mg^2+^ concentrations in the buffer or enzyme preparation (Fig. S4).

Based on these experimental results, we propose a biosynthetic pathway for aurantoside formation (Fig. 4b). A polyene chain is assembled from six malonyl building blocks in an iterative fashion by AtsF and chlorinated twice by AtsD and AtsE. AtsG catalyzes further extension steps with a malonyl and an Asn unit, before the thioesterase domain releases the aurantoside aglycone as a tetramic acid by Dieckmann cyclization. Subsequently, three glycosidic units are modified by AtsK, AtsL, and AtsN, and transferred by the glycosyl transferase homologs AtsH, AtsI, and AtsM. In a last step, AtsJ catalyzes the methylation of **1**, yielding congener **2**.

### The ‘*P. aureus*’ genome contains a large number of orphan BGCs

While searching the ‘P. aureus’ TSY genome for the aurantoside BGC, we unexpectedly detected numerous additional contigs with natural product genes suggested by the bioinformatics tool antiSMASH (version 5.0)^32^ and manual analysis (Fig. 4d). Based on the hybrid assembly draft, extensive gap closing experiments by combinatorial PCR and metagenomic cosmid isolation and sequencing generated a high-quality dataset of 19 additional extended biosynthetic loci that belong to diverse natural product classes (Fig. 4d). Omitted from this list are BGCs containing less than three biosynthetic domains and a phytoene synthase-encoding locus identified by antiSMASH that likely belongs to primary metabolism. All BGCs were localized in the ‘P. aureus’ core genome, except for the BGC *pau3* encoding a bimodular NRPS, which mapped to the single-contig plasmid (Fig. S2). In addition to the conventional BGC *pau1* representing a nine-module hybrid type I *cis*-AT PKS-NRPS, a large number of architecturally unusual *trans*-AT PKS and NRPS BGCs and BGC fragments were identified (*pau2*-*pau14*). Further BGCs belong to terpene (*pau15*) and ribosomally biosynthesized and posttranslationally modified peptide (RiPP) pathways (*pau16*-*pau19*). In addition to structural predictions based on tools integrated into antiSMASH, we used TransATor^33^, a recently developed bioinformatics tool for the automated prediction of polyketide structures for *trans*-AT PKSs. All predictions for substrate specificities for PKS and NRPS domains are annotated in Fig. 4d, but these analyses did not suggest a known natural product for any of the BGCs. Similarly, KnownClusterBlast analysis showed low similarities of the ‘P. aureus’ BGCs to characterized clusters, suggesting new natural product skeletons. To our knowledge, a comparable BGC richness among uncultivated bacteria has to date only been reported from ‘Entotheonella’, further adding to the remarkable similarities between these unrelated microbes inhabiting the same sponge.

### ‘Poriflexia’, a novel lineage of *Porifera*-associated filamentous *Chloroflexi*

In addition to their unusual genome size greatly exceeding that of previously reported *Chloroflexi*, ‘P. aureus’ TSY exhibits, to the best of our knowledge, a new morphology for this phylum. Most of the filamentous *Chloroflexi*, which belong to the classes *Chloroflexia, Anaerolineae*, and *Caldilinea*, are highly elongated, being longer than 100 μm and <1-2 μm in width, and their filaments bend flexibly. In contrast, ‘P. aureus’ TSY filaments are 30-100 μm in length and 5-10 μm wide, covered by a sheath, and straight. To further assess their affiliation, a phylogenetic tree based on 43 bacterial marker genes was constructed using genomes of filamentous *Chloroflexi* in the Refseq database (Fig. 5a). Concatenated sequences of 6,988 amino acid residues were generated from translated marker genes and aligned by CheckM. While most known filamentous *Chloroflexi* bacteria belong to the classes *Chloroflexia* or *Anaerolineae*, the phylogram supported a distinct, deep-branching clade for ‘*P. aureus*’. This phylogenetic dataset also suggested that members of the class *Caldilineae* are most closely related to ‘P. aureus’, in agreement with the 16S rRNA gene-based phylogenetic analysis (Fig. S5). For the new deep-rooting *Chloroflexi* lineage we propose the name ‘Poriflexia’, which seems to be different from the known Porifera-associated *Chloroflexi* (Fig. S6)^29^.

**Figure. 5:**
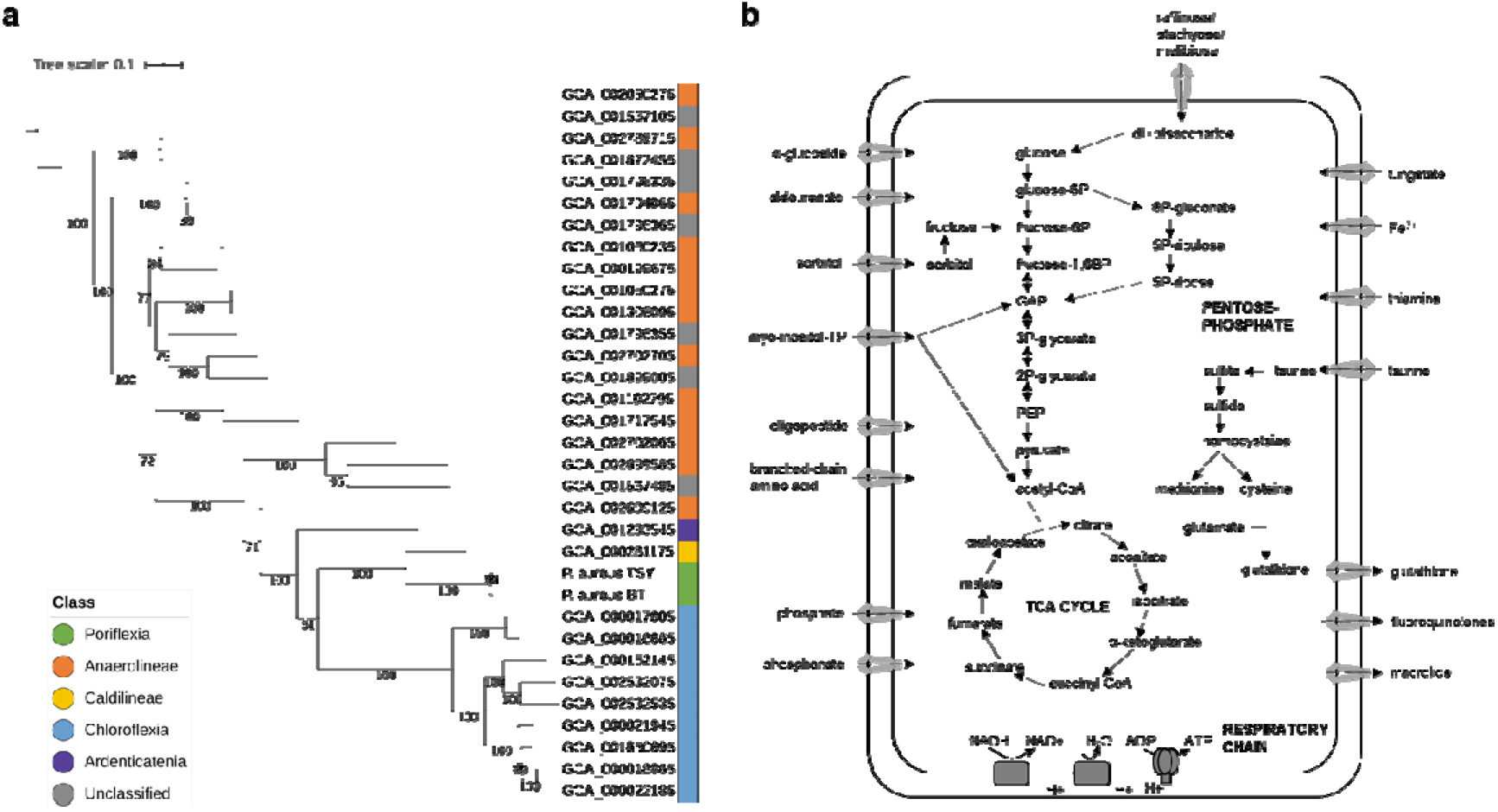
Taxonomical and functional annotation of ‘P. aureus’ TSY. **a)** The phylogram was inferred based on concatenated marker genes by 1,000-bootstrap RAxML analysis. Concatenated sequences were generated from marker genes detected in all compared Chloroflexi genomes by CheckM. Color bar beside the branch labels shows the class of the genomes. **b)** Metabolic pathways of ‘P. aureus’ TSY as suggested by the presence of genes encoding transporters.

### Metabolic pathway of ‘P. aureus’

In the draft genome of ‘P. aureus’ TSY, gene sets for glycolysis (Embden-Meyerhof-Parnas), tricarboxylic acid (TCA) cycle, the pentose phosphate pathway and a respiratory chain were found (Fig. 5b), suggesting that ‘P. aureus’ is heterotrophic. Based on analyzing preserved metabolic pathways, the symbiont seems to perform the assimilatory reduction of sulfate and lack the nitrate/nitrite reduction pathway, features resembling those of other sponge-associated *Caldilineae* bacteria^34^. ‘P. aureus’ is prototroph for vitamins B_3_ and B_6_ and auxotroph for most B-vitamins (B_1_, B_2_, B_5_, B_7_, B_9_ and B_12_), similar to its closest characterized homolog, *Caldilinea aerophila* DSM 14535 which is prototroph for B_6_ and B_12_ and auxotroph for all other B-vitamins^35^ (Table S6). For thiamine (B_1_-pathway), riboflavin (B_2_) and cobalamin (B_12_) uptake, homologs to known transporters from *Chloroflexus aurantiacus* are encoded in the ‘P. aureus’ genome^35^ (Table S7). In addition, known virulence mechanisms like type III or type VI secretion systems were not found.

An orthologue analysis using Orthofinder^36^ detected 518 core orthologous gene groups across the dataset of published filamentous *Chloroflexi* genomes (Table S4). The total number of core CDSs is 2.2 times higher in the ‘P. aureus’ TSY genome than in the other *Chloroflexi* genomes (Fig. S7). The symbiont genome contains highly amplified CDSs annotated in the COG database as “amino acid transport and metabolism”, “carbohydrate transport and metabolism”, “inorganic ion transport and metabolism”, and, as highlighted above, “secondary metabolites biosynthesis, transport and catabolism”. The amplification ratios were from 3.29 to 6.20 (Fig. S7), suggesting a strong bias to transport and metabolism function. The gene enrichment pattern of ‘P. aureus’ BT resembles that of ‘P. aureus’ TSY. Further investigation of protein functions by InterProScan showed that 8,140 CDSs in the orthologous gene groups of ‘P. aureus’ TSY are associated with 2,229 Pfam families (Table S8). Of these, 158 families were enriched in the symbiont (p-value < 0.01) compared with other *Chloroflexi* genomes, and the enrichment patterns were more similar to other sponge symbionts rather than *Chloroflexi* spp. (Fig. S8). Among the protein families, an enrichment in 775 ATP-binding cassette (ABC) transporter proteins is striking. The functional classification of ABC transporter proteins was visualized by SimGraph, which clustered the putative orthologous CDSs. About 50% of ABC transporter proteins were found to form ten large clusters (Fig. 6a). The sizes of the top five clusters were extremely large compared with SimGraph plots on other bacteria (Fig. 6b). Most of the transporter proteins were annotated as components of peptide uptake (clusters 1, 2, 4, 9, and 10), sugar uptake(clusters 3 and 5), or ligand export (clusters 6, 7, and 8) systems. This transporter enrichment pattern might be a consequence of a symbiotic lifestyle involving the acquisition of host metabolites and export of bioactive natural products. Note that the peptide importer clusters contain many glutathione importers (GsiA, C, D) although the glutathione biosynthetic pathway was detected only in the ‘P. aureus’ TSY genome among the filamentous *Chloroflexi* genomes. These facts suggest that glutathione should have a yet-to-be-uncovered important role in the ‘Poriflexus’ survival. The highly duplicated ligand multidrug transport systems (YheI, MsbA) might defend ‘Poriflexus’ from various secondary metabolites including those derived from the ‘Entotheonella’ variants in the sponge holobiont, in addition to exporting its own bioactive substances.

**Figure. 6:**
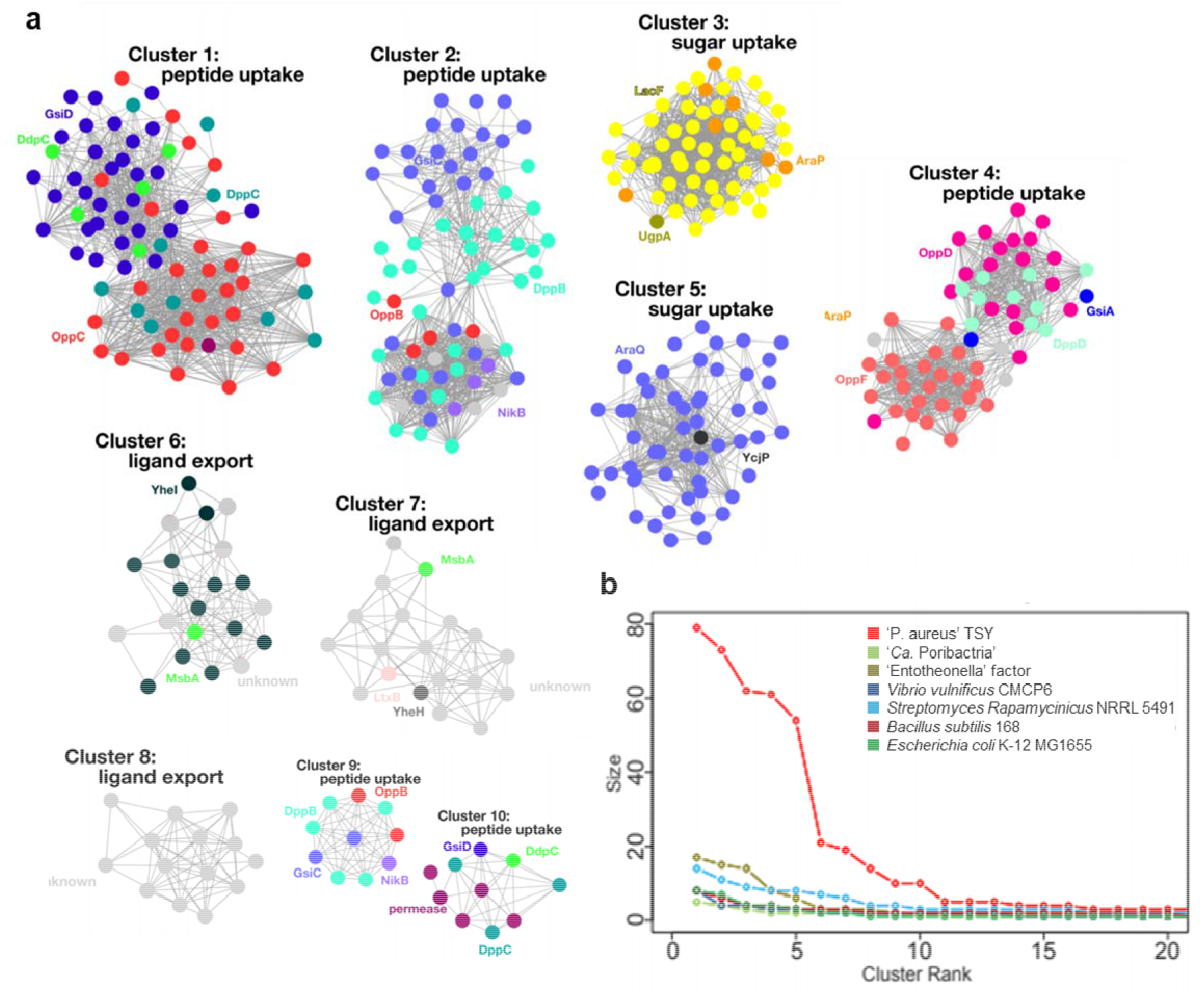
SimGraph analysis of ABC transporter proteins of ‘P. aureus’ TSY. **a)** Orthologous transporter proteins (fasta36 E-value < 10^−10^) were clustered. Colored dots show each transporter protein and its annotated function. **b)** The size distribution of ABC transporter clusters is shown against the cluster size rank in different bacteria.

## Discussion

In this study, we present the MERMAID analysis pipeline that combines Raman microscopy-based screening with single-cell analysis to link metabolic with genomic features in uncultivated bacteria. Raman microscopy is one of the few single-cell techniques that provides a direct link between metabolites and source organism in a non-destructive way and at the single-cell level and can be applied to diverse questions in microbiological research using commercially available instrumentation. Raman microscopy was previously applied to single-bacterial natural product imaging^37^ and to bacterial sorting based on deuterium incorporation^38^, but to our knowledge MERMAID is the first single-cell pipeline that also provides genomic information. This pipeline can be a one-stop, high-throughput system based on microfluidics. Recently, a single-cell analysis method linking metabolic and genomic features has been developed based on labeling of synth(et)ase enzymes^39^. The potential advantage of MERMAID is the detection of various types of intracellular biomolecules with specific molecular features. Some Raman-active moieties that do not overlap with those of basic metabolites are conjugated olefins, alkynes, carbon-halogen bonds, heteroaromatic rings, i.e., features present in many secondary metabolites. In some cases, compounds with more common features can also be targeted using analysis with MCR-ALS^37^. As an example for its utility, the platform enabled the rapid identification of the unusual symbiont ‘P. aureus’, a key provider of bioactive substances in microbiomes of two different *Theonella* sponges. The organism had remained obscure due to its morphological resemblance to ‘Entotheonella’ and low genome coverage in previous metagenomic sequencing rounds, emphasizing the importance of complementary technologies to obtain insights into microbiome functions.

‘P. aureus’, a member of a new deep-branching *Chloroflexi* lineage, shares several noteworthy features with the distantly related ‘Entotheonella’, including a large genome size, filamentous morphology, and a high number of natural product BGCs suggesting a rich specialized metabolism. BGC richness is relatively uncommon among bacterial lineages and has been reported for only a few organisms from microbial dark matter^8,40,41^. The genome sizes of ‘P. aureus’ and ‘Entotheonella’ members exceed those of all other previously reported >1200 sponge-associated bacteria^29^ by at least 3 Mb in size. The remarkable presence of two distinct non-classical producers with expanded genomes and specialized metabolism in the same sponge might be due to selective mechanisms that foster interactions with chemically rich bacteria. Among the benefits that a diversified chemistry could provide to the sessile animals are host defense against predators and epibionts, killing of eukaryotic prey, and in the case of aurantosides, protection against fungal pathogens.

*Chloroflexi* are one of the most common phyla present in sponge microbiomes^34^. The phylum includes metabolically highly diverse members such as heterotrophs and non-oxygenic photosynthetic autotrophs, aerobes and anaerobes, thermophiles, halophiles, and dehalogenating respirators. Since the cultivated representatives cover a comparably low phylogenetic diversity of *Chloroflexi* and the accumulated genomic information is insufficient, many gene functions in the ‘P. aureus’ draft genomes are still unknown. On the other hand, genome-wide analysis detected more BGC candidates than expected besides the aurantoside cluster. This high chemical potential of ‘P. aureus’ is similar to ‘Entotheonella’ and suggests a hidden natural product diversity that is much larger than the already remarkably rich described chemistry of the *T. swinhoei* holobiont. Moreover, gene enrichment analysis of the ‘P. aureus’ TSY genome revealed numerous ABC transporter genes and other functional genes commonly enriched in uncultured sponge-associated bacteria like ‘Entotheonella’ or ‘Poribacteria’, suggesting further functional commonalities between these symbionts.

## Materials and Methods

### Sponge collection

Specimens of the marine sponge *Theonella swinhoei* Y (“Y” refers to the chemotype with yellow interior containing onnamides, polytheonamides, and aurantosides) were collected at a depth of 15 m into a bucket filled with seawater around Hachijo-jima island (33°8′16″N, 139°44′5″E) in Tokyo, Japan in November 2017. The blue *Theonella* sp. was collected at a depth of 12 m from the west side of Shimoji island (24°49’24”N, 125°08’07”E) in Okinawa, Japan in October 2018. The collected sponges were cut into pieces and stored at 4 °C in Ca- and Mg-free artificial sea water (CMF-ASW)^42^.

### Preparation of bacterial fractions

To prepare filamentous bacterial fractions, an adjusted protocol based on a previously reported method^8^ was used. A 1 cm-square block of sponge tissue stored in CMF-ASW was minced in a 25 ml tube (IWAKI) by a surgical knife and incubated for 15 minutes in 10 ml of CMF-ASW on ice. During the incubation, the mixture was stirred every 5 minutes to suspend bacteria. Then, the mixture was passed through a 40 μm mesh, and the flow-through was collected in a 15 ml tube. The collected flow-through was centrifuged at 1,000 ×g for 10 minutes to pellet filamentous bacteria. Supernatants including unicellular organisms were removed. After resuspension of the pellet in 5 ml CMF-ASW, steps from the centrifugation to the resuspension were repeated several times. The mixture was allowed to settle for 30 seconds and supernatants were transferred to a clean 15 ml tube twice to remove dense sponge tissue. The enriched filamentous bacterial fraction was pelleted again and resuspended in 2 ml of CMF-ASW.

### Isolation of single filamentous bacteria

The storage medium of the filamentous bacterial fraction was replaced by Dulbecco’s Phosphate Buffered Saline liquid (DPBS, Thermo Fisher Scientific) with 1.5% ultra-low gelling temperature agarose A5030 (Sigma-Aldrich). To reduce the risk of contamination introduced by manual picking of single filaments, bacteria were encapsulated into microdroplets at a concentration of 0.1 particle/droplet by a previously fabricated microfluidic device^19^. The cell suspension with 1.5 × 10^3^ particles/μL and 2% Pico-Surf™ 1 in Novec™ 7500 (Dolomite) as carrier oil were loaded to the microchannel through polytetrafluoroethylene (PTFE) tubing (AWG24) by a Mitos P-pump (Dolomite), and droplets (50 µm in diameter) were generated. Collected droplets were solidified on ice and moved into the water phase from the oil phase as previously reported^43^. Droplets containing single filaments were manually picked using a micropipette (Drumond) under microscopic observation and isolated into a 384-well glass bottom (Corning) plate with 1.5 µl of DPBS.

### Raman spectroscopy

All Raman spectroscopic measurements were performed by using a laboratory-built 532-nm-excited Raman microspectrometer, as described in our previous report^23^. With a confocal setup, by a 100×, 1.4-NA objective lens both for laser excitation and scattered light collection, the spatial resolution of Raman imaging system was 0.3 × 0.3 μm in the lateral directions and 2.6 μm in the axial direction. The spectral resolution was 3.0 cm^-1^. For measuring the standard spectra, a solution of aurantoside A in DMSO was prepared. The solution was placed between two coverslips, and sealed with nail polish. The laser power and exposure time were set to 15 mW and 60 seconds, respectively. For the Raman imaging measurements of filamentous bacteria, 1 μL bacterial suspension was also placed between two clean coverslips, and sealed with Vaseline. By controlling the amount of suspension medium, it was possible to physically immobilize the bacteria. In the mapping measurements, the sample stage was automatically translated with 0.5 μm interval, and the exposure time was 1 second for each measured point. For the screening of aurantoside-producing bacteria, each filament bacterium isolated in 384-well plate was measured at single-cell resolution with 1 second exposure time. In the measurement of bacteria, the laser power was set 0.5 – 5 mW.

In the data preprocessing, the obtained Raman spectral data were wavenumber-calibrated, intensity-corrected, and noise-reduced by singular value decomposition analysis (SVD), using IGOR Pro software (WaveMetrics, Inc., Lake Oswego, OR, USA). Following the data preprocessing, a multivariate analysis, namely multivariate curve resolution-alternating least squares (MCR-ALS) was conducted based on the method developed in our previous study^20^. In this study, all the spectral data obtained from bacterial mapping measurements were combined into one matrix, and the non-negative matrix factorization was performed with the SVD initialization method and L1-norm regularized ALS optimization method. All the MCR-ALS calculations were performed by an in-house program written inby python, using the SciPy library (https://github.com/mshrAndo/PyMCR). For the screening of aurantoside producing bacteria, the MCR-ALS decomposed spectra were used as reference spectra, and the presence of aurantosides was detected by least-squares spectral fitting.

### Whole-genome amplification of single filaments

The Raman microscopy-selected bacteria were lysed by incubation at 37 °C for 30 minutes with Ready-lyse lysozyme (Epicentre) of 10 U/µl and heat treatment at 95 °C for 10 minutes. To acquire single-amplified genomes (SAG), multiple displacement amplification (MDA) reaction mix (REPLI-g Single Cell Kit, QIAGEN) was added to the lysate and incubated at 30 °C for 3 hours, and the amplification was terminated by incubation at 65 °C for 3 minutes. As compared to the manufacturer’s protocol, the volume of the MDA reactions was scaled down to 10 µl to amplify genomes in the 384-well plate. MDA products were assessed by PCR and sequencing of 16S rRNA gene fragments (V3-V4 region).

### Whole-genome sequencing

Illumina sequencing libraries were prepared from each MDA product mixture using the Nextera XT kit. Libraries were sequenced in a 2 × 301-bp paired-end run by MiSeq (Illumina), and 12 SAG data of ‘P. aureus’ with a total size of 5.26-Gb. To evaluate the acquired draft genomes, metagenomic shotgun sequencing was also conducted using a procedure carefully adapted to lyse ‘Entotheonella’ filaments to enrich ‘P. aureus’ genomic DNA. Metagenomic DNA was extracted by DNA Clean & Concentrator Kit (ZYMO research) from lysate of the filamentous bacterial fraction acquired by incubation for 8 hours with 10 U/µl Ready-lyse lysozyme (Epicentre). The sequencing library was prepared using the Nextera XT kit and sequenced the same way as for the SAGs.

### Metagenomic cosmid library screening

To isolate cosmids harboring PKS genes, a metagenomic cosmid library, previously^44^ prepared from *T. swinhoei* Y total DNA in the pWEB vector (Epicentre), was screened and a 3D growth-and-dilution library format as reported earlier^45^. The isolated cosmids pTSTA2 and pTSTA3 were sequenced using Sanger technology.

### Biochemical assays with the aurantoside methyltransferase AtsJ

Based on the *atsJ* sequence, a codon-optimized gene for heterologous gene expression in *E. coli* was synthesized (Integrated DNA Technologies). Flanking restriction sites for *Nde*I and *Eco*RI were added, resulting in the plasmid pUCIDT_*atsJ*. The *Nde*I and *Eco*RI-excised insert of pUCIDT_*atsJ* was ligated into the same restriction sites of pET-28b (Novagen) and the resulting plasmid was introduced into *E. coli* DH5 α by electroporation. The construct was confirmed by DNA sequencing.

For protein expression, electrocompetent *E. coli* Tuner™ (DE3) cells (Novagen) were co-transformed with the plasmids pET-28b_*atsJ* and pG-KJE8 (TAKARA). A 200 ml culture in TB medium (50 μg/ml kanamycin, 20 μg/ml chloramphenicol, 0.5 mg/ml L-arabinose, and 5 ng/ml tetracycline) was grown at 37 °C and 180 rpm to an OD_600_ of 0.5 - 0.8, cooled on ice for 5 minutes, induced with isopropyl β-D-1-thiogalactopyranoside (0.1 mM final concentration), then shaken for 20 hours (180 rpm, 16 °C). Cells were collected by centrifugation and homogenized in 1 mL per 100 mg dry cell material in assay buffer (AB, 50 mM phosphate buffer, pH 8.0, 300 mM NaCl, 10% [v/v] glycerol). The suspension was placed on ice, and cells were lysed by sonication. Cell debris was removed by centrifugation at 12,000 ×g at 4 °C for 20 minutes and the cleared supernatant was either directly used for the reconstitution assays or subjected to purification with Ni-NTA agarose (Macherey-Nagel). The suspension was incubated for 60 minutes and transferred to a fretted column. The resin was washed with AB with 20 mM imidazole and eluted with 3×0.5 ml elution buffer (AB with 250 mM imidazole). Subsequently, imidazole was removed using a PD-minitrap column (GE Healthcare) for buffer exchange to AB.

The assays were conducted in a total volume of 200 μl of AtsJ in AB, supplemented with 5 mM MgCl_2_, 1 mM SAM and 0.25 mM aurantoside B. Negative controls with concentrated lysate from *E. coli* Tuner™ (DE3) pG-KJE8 pET28b(+), boiled concentrated lysate of *E. coli* Tuner™ (DE3) pG-KJE8 pET28b(+)_*atsJ* (10 min at 98 °C) and without addition of SAM were analyzed in parallel. After incubation for 3 hours at 30 °C and 180 rpm in the dark, samples were prepared for ultra-high performance liquid chromatography-high resolution heated electrospray-mass spectrometry (UPLC HR HESI MS) analysis in the dark using ZipTip^®^C18 (Millipore) purification and resuspension in acetonitrile.

The UPLC HR HESI MS experiments were performed on a Dionex Ultimate 3000 UHPLC coupled to a Thermo Scientific Q Exactive™ mass spectrometer. A solvent gradient (A = water + 0.1% formic acid and B = acetonitrile + 0.1% formic acid with 95:5 A/B ramped to 100% B over 10 min, 100% B for 5 min, ramped to 95:5 A/B over 1 min at a flow rate of 1.0 ml/min) was used on a Kinetex C18-XB (2.6µ, 150 × 4.6 mm) column at 27 °C. The MS was operated in positive ionization mode at a scan range of 100 to 1,000 *m/z* with a scan resolution of 70,000. The spray voltage was set to 3.7 kV and the capillary temperature to 320 °C. Spectra were annotated using the software Xcalibur™ version 4.1 (Thermo Fisher Scientific).

### Assembly and annotation of the ‘Poriflexus aureus’ genome

The quality of each SAG dataset was controlled by removing low-quality reads (≥50% of bases with quality scores□<□25), trimming the 3′-end low-quality bases (quality score□<□20), and removing <20 bp reads and reads including >1% unidentified bases. Contigs were assembled from each SAG dataset by SPAdes 3.12.0^24^ with “--sc -- careful” option. Then, a composite draft genome was generated by ccSAG^25^. The assembly quality of composite draft genomes and drafts from each SAG data were evaluated by QUAST 4.5^26^ and genome completeness and contamination rate of them were assessed by CheckM 1.0.6^27^.

Gene annotation and metabolism prediction of the composite draft genomes was conducted by Prokka 1.13^28^, Genomaple^46^, and BLAST search^47^ to COG database^48^. All high-quality *Chloroflexi* genomes (>90% genome completeness, <10% contamination rate) except for the class Dehalococcoidia composed of unicellular *Chloroflexi* sp. and genomes of five sponge-associated bacteria (two ‘Entotheonella’ and three ‘Poribacteria’ variants) used in the comparative genome analysis were downloaded from the NCBI genome database (Table S4) and annotated by the same protocol as for the composite draft genomes. *Chloroflexi* phylogenetic trees were inferred by the maximum likelihood method using RAxML v 0.9.0^49^ based on concatenated marker genes identified by CheckM or 16S rRNA genes. Functional enrichment analysis of the ‘Poriflexus’ genomes were conducted by Fisher’s exact test using Pfam annotation result, and the result was visualized by iTOL^50^. Orthofinder^36^ was used for finding orthologous gene groups encoded in the different genome sequences and to estimate the degree of gene duplication in each genome sequence. SimGraph, an in-house tool for visualizing gene duplication based on fasta36^51^ and Cytoscape 3.8.0^52^, was used to compare the degree of duplication of a specific gene. Briefly, all-against-all similarity calculation was carried out among the amino acid sequences derived from the duplicated genes by fasta36 and pairs with E-value < 10^−10^ were connected with edges on Cytoscape.

### Long-read sequencing and analysis of BGCs

To acquire continuous sequences of BGCs, long-read sequencing was conducted. A sequencing library was prepared by Rapid Sequencing kit (Nanopore) from a mixture of five single-filament MDA products generating high-quality SAG short reads and sequenced by MinION (Nanopore) for 48 hours. From the acquired 1.48-Gb long reads, long-read contigs were assembled by Canu 1.4^53^ and polished by Pilon^54^ using short-read sequences of 12 short-read SAG samples. Gene annotation of long-read contigs was conducted by Prokka and BGCs were identified and annotated by antiSMASH 4.0.2^55^. Predictions of natural product structures for *trans*-AT PKSs were generated by TransATor^33^. Manual BLAST analyses for verification of BGCs and discovering of undetected BGCs were carried out in addition.

## Data availability

Sequence reads and genomic sequences of ‘P. aureus’ have been submitted to the National Center for Biotechnology Information Search database under project number PRJNA681468. Individual accession numbers of compared bacterial genomes can be accessed in Supplementary Table 3. The aurantoside BGC is registered to the MiBIG (ID BGC0002131).

## Acknowledgments

This work was supported by MEXT KAKENHI grant numbers 17H06158 to H.T., by the Gordon and Betty Moore Foundation (grant GBMF9204, DOI: https://doi.org/10.37807/GBMF9204) and the Swiss National Science Foundation (205320_185077) to J.P. We thank C. Sakanashi, H. Maciejewska-Rodrigues and H. Minas for technical support. The super-computing resource was provided by the Human Genome Center (University of Tokyo).

