## Supplementary material for "Single-cell metabolite detection and genomics reveals uncultivated talented producer": Supplemantal Document

Supplementary Information Text

Evaluation of the composite ‘P. aureus’ TSY draft genome

The ‘P. aureus’ TSY genome of 14 Mbp length, as suggested by the combined SAG data, was re-assessed by three different methods because of the unusually large genome size compared to other *Chloroflexi* bacteria (4.5 Mbp on average and up to 8.1 Mbp). Firstly, the estimated genome size of ‘P. aureus’ TSY was calculated from each SAGs based on the total contig length and the estimated genome completeness. The estimated genome size converged to around 13.5 Mbp (Fig. S1a). Second, contigs in the draft genome were binned by mapping results of metagenomic shotgun sequencing reads to the contigs (Fig. S1b). Metagenomic short reads derived from the filamentous fraction were mapped to contigs in the combined SAG data. Contigs were binned based on GC content and sequence depth, and most of the contigs gathered into one cluster. Clustered contigs were also annotated taxonomically with BLAST search to nr database, which confirmed that most of the genomic sequences were derived from bacteria and the draft genome contained few eucaryotic contaminations. Third, sequence appearance patterns of the contigs from each single-cell data were investigated in the following way: SAG short reads and metagenomic short reads were remapped to contigs of the draft genome, and the number of SAGs mapping to each genomic position was counted. Because most contigs were covered by >= 6 SAGs and more than 99% of contigs were covered by metagenomic data, it was confirmed that the draft genome hardly contained contamination derived from DNA amplification.

NRPS-encoding plasmid pPAUTSY01

The hybrid assembly output contained one circular contig (Fig. S2a) of 335,928 bp harboring 380 CDSs containing the NRPS BGC *pau3*. The CDSs included also genes for ParA/ParB related to a plasmid partitioning system for low-copy plasmids^4^ and for the DNA replication initiator protein (PF10134), suggesting that the circular contig is a plasmid of ‘P. aureus’ TSY. The fact that the circular contig was a plasmid sequence was also supported by binning analysis of contigs in the Illumina composite draft genome suggesting that contigs mapping to the circular sequence were about 1.5 times as abundant as those of the core genome (Fig. S2b).

Analysis of ‘P. aureus’ in the blue *Theonella* sp. sponge

Raman spectroscopy and genome analysis of ‘P. aureus’ BT (Fig. S3a) was conducted in the same way as the experiments using ‘P. aureus’ TSY. Raman spectra obtained from filamentous bacteria isolated from the blue sponge (*Theonella* sp. BT) were analyzed by MCR-ALS analysis. As shown in Fig. S3c, decomposed spectra assignable to secondary metabolites were almost identical to the decomposed spectra of ‘P. aureus’ TSY. LC-MS analyses of extracts from the blue *Theonella* sponge also showed the existence of aurantosides as in *T. swinhoei* TSY (Fig. S3b). After Raman microspectroscopy, 7 SAGs of ‘P. aureus’ BT were sequenced and a composite draft genome with 68% genome completeness was assembled (Table S1, S2). The acquired draft genome of ‘P. aureus’ BT was compared to the draft genome of ‘P. aureus’ TSY. With a 99.5% identity of full-length 16S rRNA genes and an average nucleotide identity 98% the two symbionts can be considered the same candidate species (Fig. S3d). Furthermore, the aurantoside BGC of ‘P. aureus’ TSY was also highly conserved in ‘P. aureus’ BT (Fig. S3e). As with the Raman microscopy, these sequencing results supported aurantoside biosynthesis by ‘P. aureus’ BT.

Core genome of ‘P. aureus’ as a member of the phylum *Chloroflexi*

Ortholog analysis using CDS sequences by orthofinder was performed, and 22,531 OGs (orthogroups) were constructed from 118,007 CDS of 33 genomes of filamentous *Chloroflexi* (Table. S4). Then, the core genome size was estimated to assess the presence of *Chloroflexi* core OGs which are highly conserved across the diverse *Chloroflexi* genomes. In previous research^5^, it was reported that the number of core OGs identified from *n* genomes can be formulated by the empirical determining functions

Fc (n)= κc exp[-n/τc ]+Ω

(Ω: the asymptotic core genome size, κc: the magnitude of the decay, τc: the rate at which the curve approaches Ω)

Permutation tests of strict and extended core OGs, which include at least one CDS from every *n* genome or at least *n-1* genomes, were conducted (Fig. S7a). The strict core genome size Ωc and the extended core genome size Ωe were estimated as 290 and 500 by fitting the formula to the permuted core OGs, and presence of the *Chloroflexi* core OGs was indicated. The 22,531 OGs from the 33 *Chloroflexi* genomes were classified as 518 extended core OGs, 11,559 unique (species-specific) OGs, and 10,454 character OGs (remains) (Fig. S7b). In pan genome analyses comparing genomes of closely related species, it has been reported that U-shaped frequency distributions are obtained because most OGs are assigned as core or unique OGs^6^. In this study, although more diverse genomes across the phylum *Chloroflexi* were compared than those in the typical species- or genus-level pangenomic analysis, it was confirmed that the frequency distribution follows a gradual U-shape. Focusing on the ‘P. aureus’ TSY genome, the ratio of CDSs in the extended core OGs was 23.4%, which is slightly lower than other *Chloroflexi* genomes (26.2-40.6%) because of many CDS in unique OGs, but the number of CDSs in the extended core OGs was 2,755 and about 2.2 fold that of the other *Chloroflexi* genomes (1,009-1,807, average: 1,260) (Fig. S7c). Then, enriched functional CDSs were investigated from the COG annotation result by Prokka. The ‘P. aureus’ genomes had highly amplified CDSs annotated in the COG database as "amino acid transport and metabolism", "carbohydrate transport and metabolism", "inorganic ion transport and metabolism", and, as highlighted above, "secondary metabolites biosynthesis, transport and catabolism" (Fig. S7d).

Core and character OGs were reconstructed using genomes of *Chloroflexi* species, other sponge symbionts (‘Entotheonella’, ‘Poribacteria’) and 10 bacterial species including selected marine and various well-studied bacteria (Table S4). Then, Pfam annotation of CDS in each OG was conducted by InterProScan. The Pfam enrichment patterns of the ‘P. aureus’ genomes were more similar to those of sponge symbionts rather than those of other *Chloroflexi* genomes (Fig. S8). For example, phytanoyl CoA dioxygenase (PhyH, PF05721), sulfatase (PF00884) and pentapeptide repeats (PF00805) are to be noticed as common enriched Pfams among sponge symbionts including ‘P. aureus’. Genes encoding PhyH homologs were suggested as genes for 2-aminoethylphosphonate utilization in sponge phospholipids to provide phosphorous sources^7^. An enrichment of arylsulfatase genes was reported in diverse sponge-associated bacteria such as *Alphaproteobacteria* and proposed to utilize sulfated polysaccharides of host sponges^8^. These common features among sponge symbionts indicate the adaptation of ‘P. aureus’ to utilize sponge tissue.

### Supplementary Figures

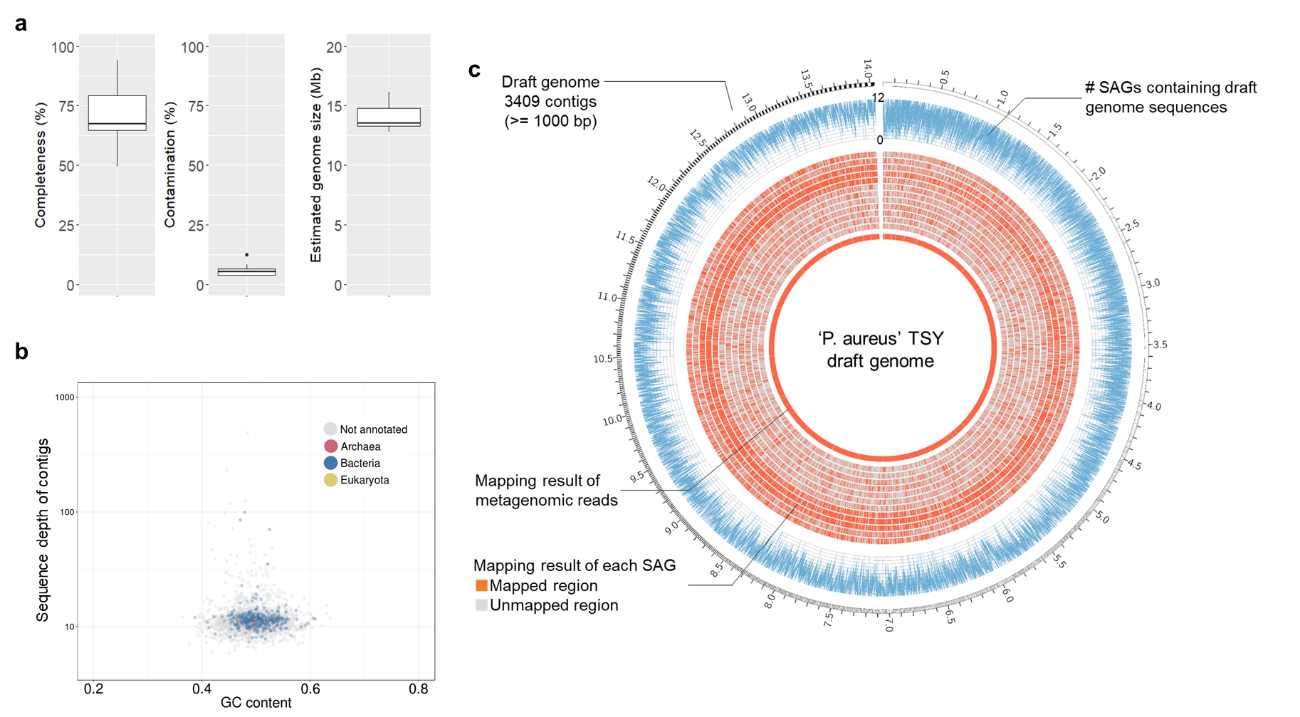

**Fig. S1:** Evaluation of the composite ‘P. aureus’ TSY genome. **a)** Individual genome size estimation from each SAG using completeness, contamination and total contig length. Box plot showing median, interquartile range, minimum and maximum values. Outlying values are shown as circles. **b)** Binning of the contigs in the ‘P. aureus’ TSY draft genome using metagenomic shotgun sequencing data. Colored points show the taxonomic annotation of the contigs. **c)** Sequence appearance pattern from each SAG. The external circle shows the total contig length of the concatenated contigs and the orange parts of internal circles show contig sequences appearing in SAGs or metagenomic data. The number of SAGs covering each genomic position is shown by the blue line graph.

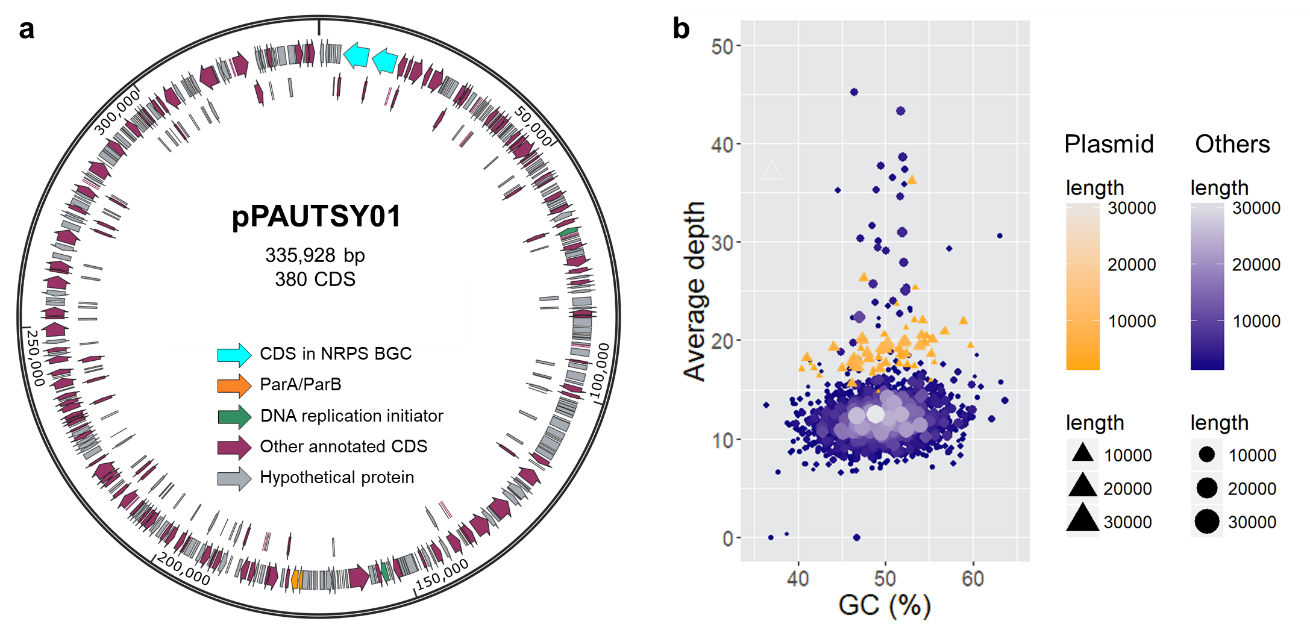

**Fig. S2:** NRPS-encoding plasmid pPAUTSY01. **a)** Gene map of the plasmid sequence containing the NRPS BGC *pau3*. The colored arrows show the annotation result of each CDS. **b)** Binning of the SAG composite draft genome. Contigs in the draft genome were binned based on GC content and sequence depth calculated from metagenomic sequencing data. The orange triangles show contigs mapped to the pPAUTSY01, and the blue circles show contigs of the core genome. Saturation and size of each point refer to the contig length.

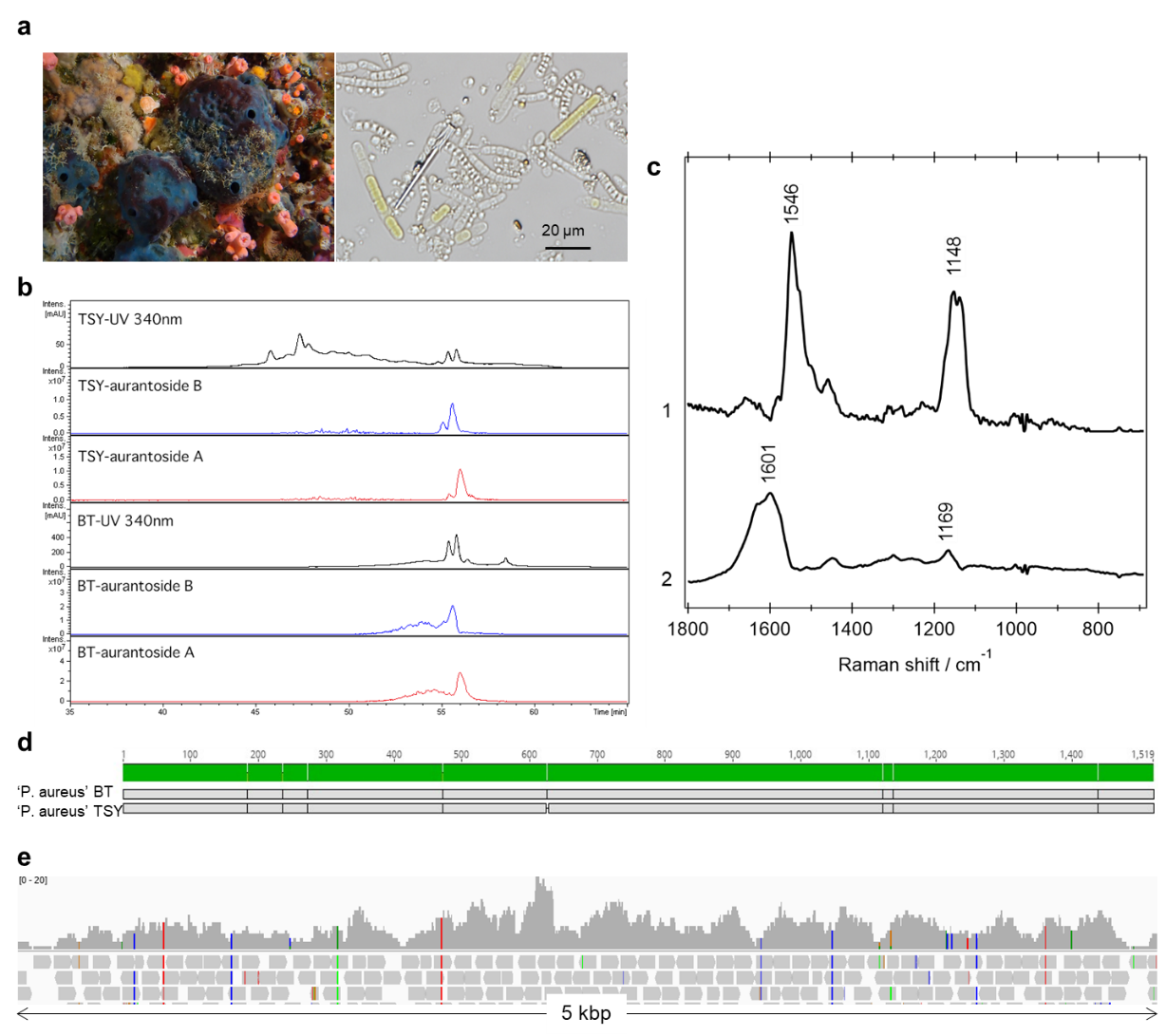

**Fig. S3:** Raman spectroscopy and genome analysis of ‘P. aureus’ BT **a)** The blue sponge *Theonella* sp. BT (left) and light micrograph of associated bacteria. **b)** LC-MS data for extracts of the sponges TSY and BT. For aurantosides A and B, extracted ion chromatograms are shown at *m*/*z* 817.4 and 803.4, respectively. **c)** MCR-ALS-decomposed Raman spectra of ‘P. aureus’ BT (component 1: aurantoside, 2: aurantoside derivative with shorter polyene length). **d)** Alignment of the full-length 16S rRNA genes of ‘P. aureus’ TSY and ‘P. aureus’ BT. Identical regions are shown in green. **e)** Aurantoside BGC locus of SAGs of ‘P. aureus’ BT. Sequence reads of ‘P. aureus’ BT were mapped to the aurantoside BGC of ‘P. aureus’ TSY, and Integrative Genome Viewer imaged the result. The upper bar chart shows the sequence depth of the aurantoside BGC in ‘P. aureus’ BT SAGs and the color shows the mapped nucleotide (gray: the same as the reference, others: different nucleotides from the reference (blue: C, green: A, red: T, orange, G)).

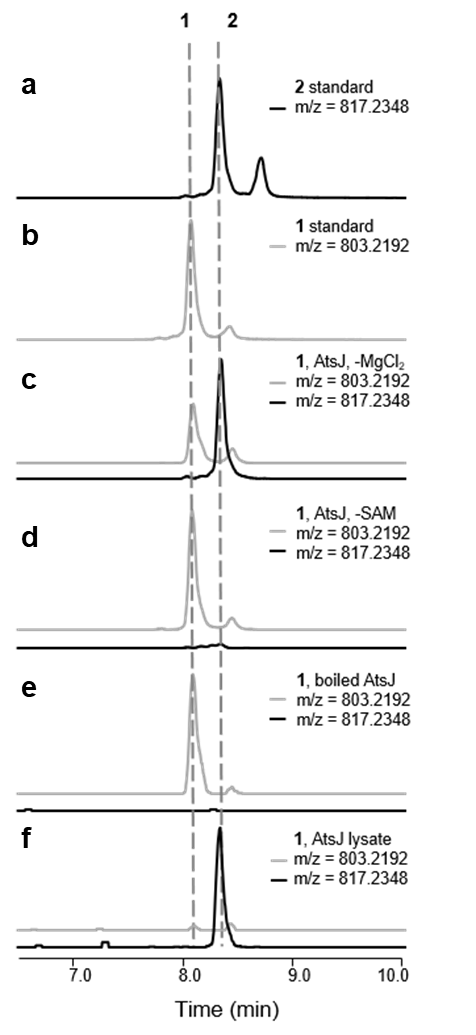

**Fig. S4:** Control experiments for reconstitution assays with the *O*-methyl transferase AtsJ (supplement to Fig. 4c). Shown are extracted ion chromatograms. **a)** Aurantoside A (**2**) standard, **b)** Aurantoside B (**1**) standard. **c)** In reconstitution assays with AtsJ without MgCl_2_, **1** is partially converted to **2**. **d)** The assay lacking *S*-adenosylmethionine (SAM) shows no conversion of **1** to **2**. **e)** The boiled AtsJ enzyme shows no conversion of **1** to **2**. **f)** Conversion of **1** to **2** is observed for the *E. coli* lysate containing AtsJ.

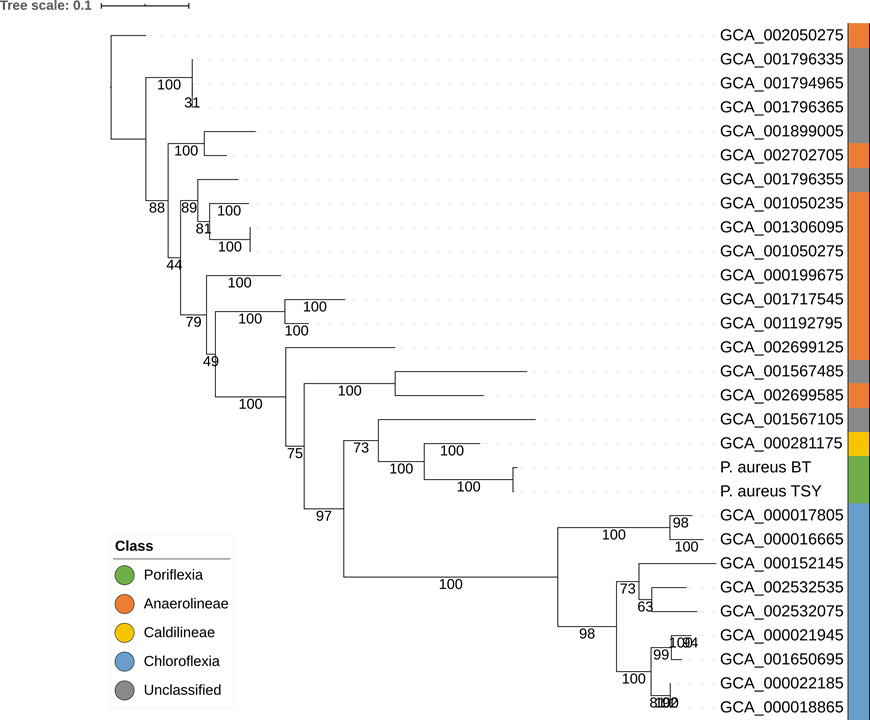

**Fig. S5:** Phylogenetic analysis of filamentous *Chloroflexi* based on whole 16S rRNA genes. The phylogram was inferred using RAxML^1^ analysis with 1,000 bootstraps.

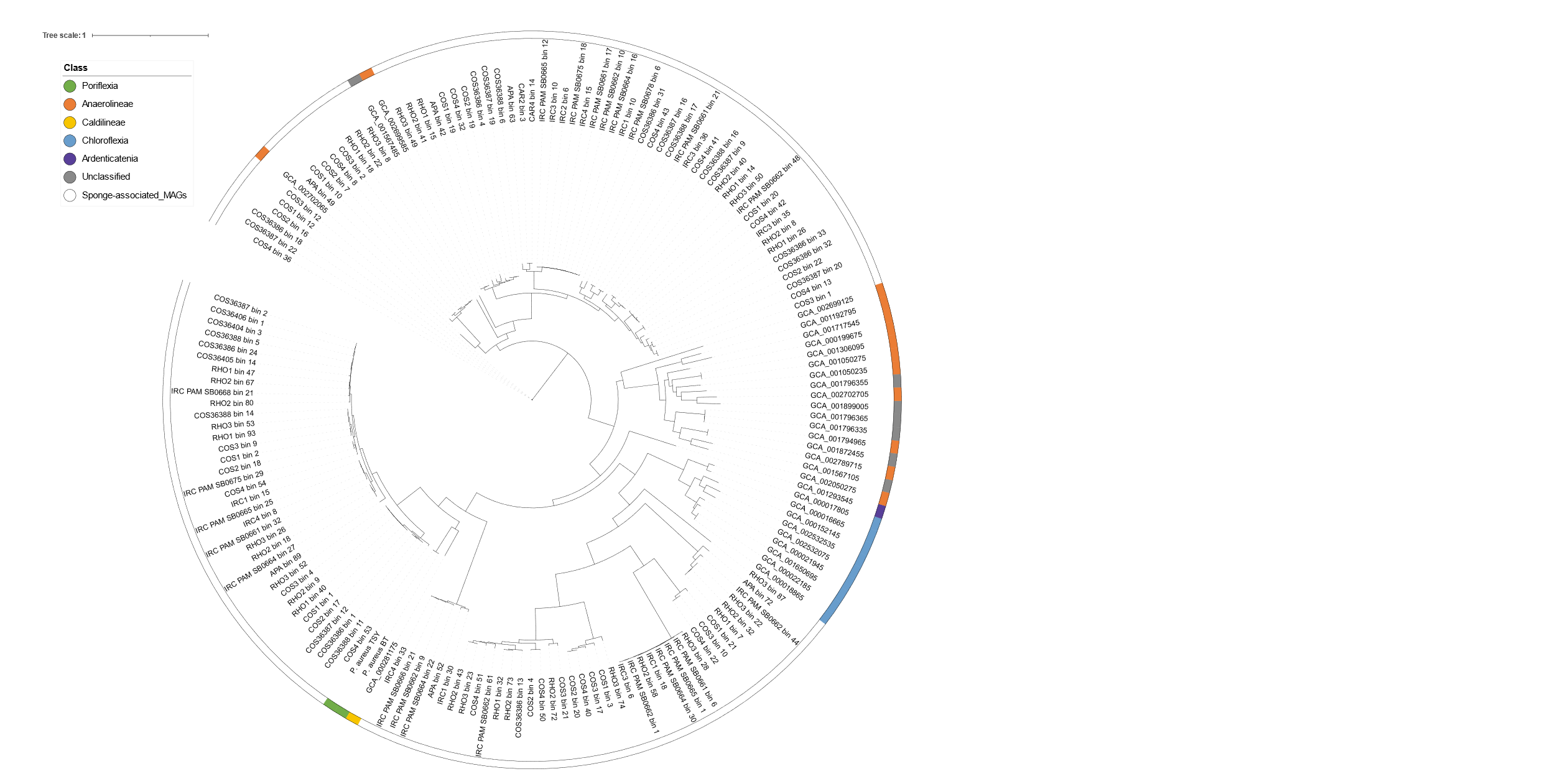

**Fig. S6:** Phylogenetic analysis of *Chloroflexi* based on concatenated marker genes. Concatenated sequences of all compared *Chloroflexi* genomes including 137 MAGs^2^ acquired from sponge-associated microbiota were generated from marker genes detected by CheckM. Color bar beside the branch labels shows the attribute of the genomes.

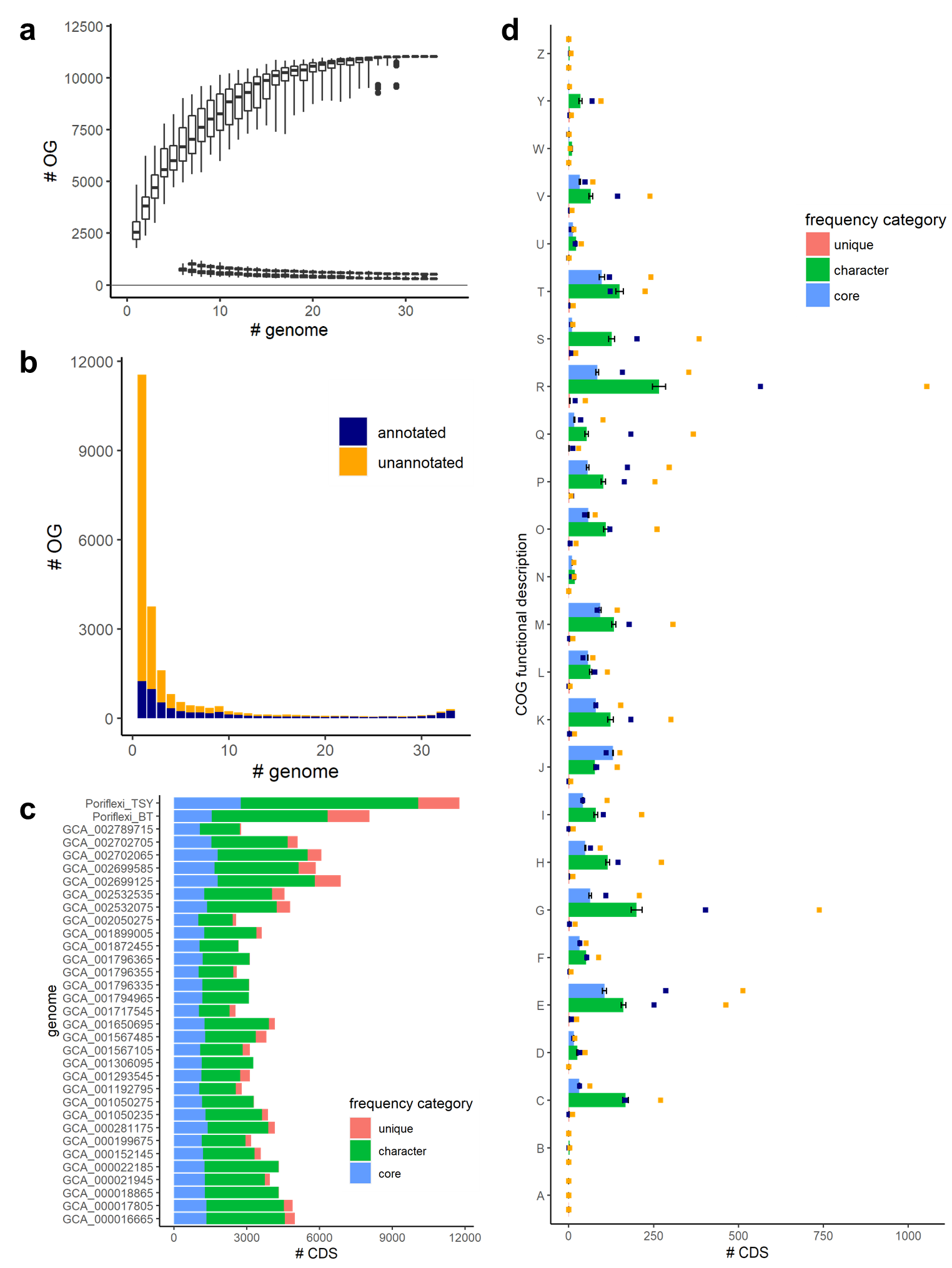

**Fig. S7:** Gene frequency in *Chloroflexi* genomes. **a)** Core genome size of 33 filamentous *Chloroflexi* including ‘P. aureus’. Identification of core OGs (orthogroups) was permuted 100 times. The upper, middle, and bottom series of box plots show the number of total OGs, the extended core OGs (including CDS from at least *n-1* genomes), and the strict core OGs (including CDS from every genome). The box plots show medians and quartiles. **b)** Frequency distribution of homology groups. The OGs constructed from 33 genomes were classified by the number of genomes composing the OGs. Navy blue shows the OGs including at least one annotated CDS, and orange shows the OGs including only unannotated CDS that were assigned by only Prokka as “hypothetical proteins”. **c)** Frequency distribution of homology groups. Number of CDS in core OGs (blue), character OGs (green), unique OGs (red) is shown per genome. **d)** Genomic functional composition of ‘P. aureus’. COG functional categories were annotated to each CDS, and the functional annotation was correlated to conserved OGs. Orange and navy plots show the number of CDS in ‘P. aureus’ TSY and ‘P. aureus’ BT, respectively. Bars shows the average of the number of CDS in the other *Chloroflexi* genomes.

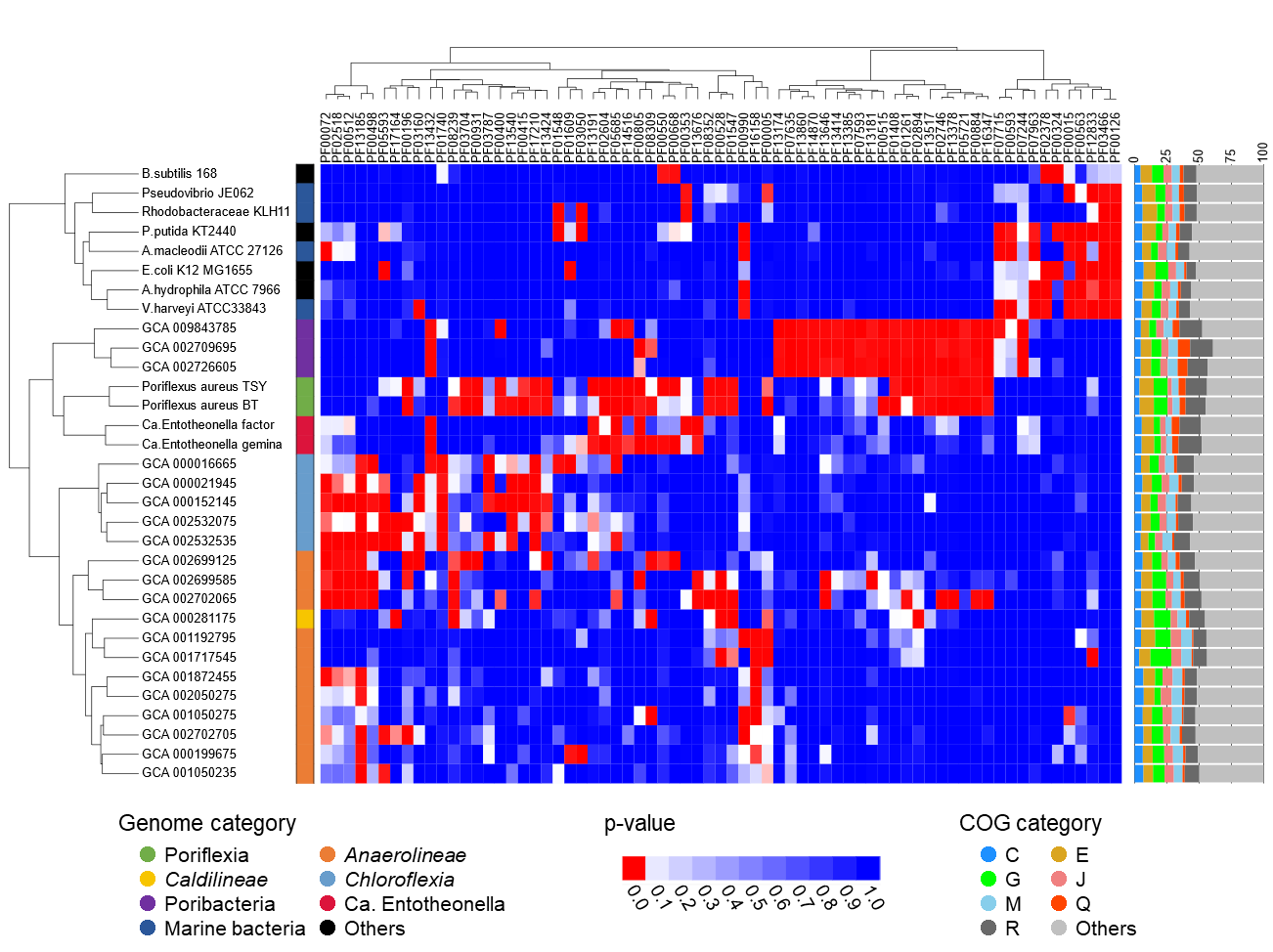

**Fig. S8:** Enrichment pattern of functional genes in core and character OGs. Genomes of filamentous *Chloroflexi*, other sponge symbionts (‘Entotheonella’, ‘Poribacteria’) and 10 bacterial species including selected marine and various well-studied bacteria were compared. Enriched Pfams in each genome were detected by Fisher’s exact test, and a heatmap was drawn based on the p-value. Genomes or Pfams which showed similar enrichment patterns were clustered by the ward method using the p-value. The left color bar shows the taxonomy of compared genomes, and the right bar chart shows the color-coded COG functional description of each genome.

### Supplementary Tables

**Table S1:** Quality of sequencing data of individual ‘P. aureus’ TSY SAGs. Samples marked with 'Y' were obtained from the *Theonella swinhoei* chemotype Y and samples marked with 'B' were obtained from the blue *Theonella* species.

| **Sample** | **Raw reads**  **(Mbp)** | **# Contigs**  **(>=1,000 bp)** | **Total length**  **(>=1,000 bp)** | **Total #**  **contigs** | **Largest**  **contig** | **GC (%)** | **N50 (bp)** | **Completeness**  **(%)** | **Contamination**  **(%)** |
| --- | --- | --- | --- | --- | --- | --- | --- | --- | --- |
| S1 Y | 59.25308 | 2,094 | 3,635,396 | 6,569 | 10,302 | 49.02 | 1,072 | 57.89 | 3.74 |
| S2 Y | 91.163392 | 3,082 | 5,811,595 | 7,712 | 10,143 | 49.28 | 1,312 | 73.50 | 6.53 |
| S3 Y | 83.90811 | 2,813 | 5,241,514 | 7,448 | 12,306 | 49.35 | 1,248 | 72.29 | 5.56 |
| S4 Y | 95.686852 | 2,578 | 5,087,374 | 7,111 | 17,650 | 48.07 | 1,268 | 65.15 | 4.65 |
| S5 Y | 89.801434 | 2,289 | 3,794,039 | 8,474 | 10,642 | 47.94 | 961 | 54.09 | 2.02 |
| S6 Y | 92.144756 | 3,074 | 5,642,938 | 7,858 | 12,089 | 49.38 | 1,277 | 71.75 | 5.20 |
| S7 Y | 95.844202 | 3,194 | 6,007,700 | 7,796 | 14,164 | 49.42 | 1,310 | 76.58 | 6.31 |
| S8 Y | 986.35925 | 3,033 | 8,640,182 | 5,459 | 26,441 | 49.01 | 2,922 | 78.23 | 5.76 |
| S9 Y | 987.81904 | 2,687 | 7,122,809 | 5,182 | 20,590 | 49.00 | 2,592 | 67.47 | 3.74 |
| S10 Y | 957.68194 | 3,412 | 12,146,580 | 5,223 | 24,996 | 49.29 | 4,412 | 91.97 | 4.19 |
| S11 Y | 912.944862 | 3,565 | 13,557,043 | 4,963 | 25,090 | 49.20 | 4,883 | 95.00 | 6.31 |
| S12 Y | 797.67928 | 3,565 | 11,955,213 | 5,547 | 32,549 | 49.17 | 3,931 | 89.82 | 6.01 |
| S13 B | 95.467414 | 874 | 2,662,744 | 7,037 | 13,417 | 47.34 | 1,761 | 29.57 | 2.30 |
| S14 B | 71.999984 | 501 | 1,517,981 | 3,195 | 13,179 | 48.63 | 2,171 | 24.37 | 1.39 |
| S15 B | 79.586605 | 435 | 1,388,548 | 3,243 | 12,947 | 48.80 | 2,279 | 15.13 | 1.72 |
| S16 B | 76.124802 | 370 | 1,191,725 | 2,324 | 16,008 | 49.36 | 2,543 | 13.79 | 0.00 |
| S17 B | 72.619849 | 516 | 1,551,609 | 3,581 | 12,304 | 48.84 | 2,074 | 13.64 | 0.91 |
| S18 B | 67.445634 | 431 | 1,324,130 | 2,862 | 13,179 | 49.03 | 2,274 | 11.99 | 1.72 |
| S19 B | 77.995783 | 449 | 1,433,244 | 3,260 | 20,788 | 49.05 | 2,304 | 11.54 | 0.86 |

**Table S2:** SAG-combined draft genomes of ‘P. aureus’ in two *Theonella* sponges. The short-read composite assemblies were constructed from multiple Illumina sequencing data of ‘P. aureus’ SAGs by ccSAG. The hybrid assembly was constructed from Nanopore sequencing data of ‘P. aureus’, SAG mixture by canu and polished with the Illumina sequencing data.

| **Bacteria** | **Assembly** | **Completeness**  **(%)** | **Redundancy**  **(%)** | **Estimated genome**  **size (Mb)** | **# Contigs** | **N50 (bp)** | **GC (%)** | **# CDS** | **# rRNA**  **operons** | **# tRNAs** |
| --- | --- | --- | --- | --- | --- | --- | --- | --- | --- | --- |
| ‘P. aureus’ TSY | Short-read composite | 95.61 | 6.21 | 14.0 | 3,409 | 5,494 | 49.7 | 11,772 | 1 | 45 |
| ‘P. aureus’ TSY | Hybrid | 92.73 | 6.82 | 15.0 | 195 | 102,784 | 49.2 | 14,903 | 1 | 44 |
| ‘P. aureus’ BT | Short-read composite | 68.17 | 4.09 | 12.9 | 4,969 | 2,574 | 48.7 | 9,221 | 1 | 48 |

**Table S3:** tRNA genes in the ‘P. aureus’ TSY genome.

| **tRNA codons** | | |
| --- | --- | --- |
| Ala (GGC, TGC) | His (GTG) | SeC (TCA) |
| Arg (ACG, CCG, CCT, GCG) | Ile (GAT) | Ser (CGA, GCT, GGA, TGA) |
| Asp (GTC) | Leu (CAA, CAG, GAG, TAG) | Thr (CGT, GGT, TGT) |
| Cys (GCA) | Lys (CTT, TTT) | Trp (CCA) |
| Gln (CTG, TTG) | Met (CAT) | Tyr (GTA) |
| Glu (CTC, TTC) | Phe (GAA) | Val (CAC, GAC, TAC) |
| Gly (CCC, GCC, TCC) | Pro (CGG, GGG, TGG) |  |

**Table S5:** ORFs detected on the 'P. aureus' draft genome belonging to the aurantoside (*ats*) BGC and their proposed protein functions.

| **ORF** | **Protein size [aa]** | **Proposed protein function** | **Closest protein homologue [source organism]** | **amino acid identity [%]** | **GenBank**  **accession number** |
| --- | --- | --- | --- | --- | --- |
| *atsA* | 751 | ABC transporter | hypothetical protein C2W62_13485 [*Candidatus* 'Entotheonella serta'] | 51 | PON17394.1 |
| *atsB* | 786 | ABC transporter | hypothetical protein C2W62_13470 [*Candidatus* 'Entotheonella serta'] | 39 | PON17391.1 |
| *atsC* | 752 | Cation transport ATPase | cation transport ATPase [*Beggiatoa* sp. PS] | 52 | EDN70595.1 |
| *atsD* | 552 | Halogenase | FAD-dependent oxidoreductase [*Streptomyces africanus*] | 50 | WP_086558590.1 |
| *atsE* | 627 | Halogenase | hypothetical protein AU255_14460 [*Methyloprofundus sedimenti*] | 48 | OQK16290.1 |
| *atsF* | 5,777 | PKS (KS, AT, DH, KR, ACP) | Acyl transferase domain-containing protein [*Lihuaxuella thermophila*] | 44 | SEM89772.1 |
| *atsG* | 7,790 | PKS/NRPS (KS, AT, ACP, C, A, PCP, TE) | non-ribosomal peptide synthetase [*Moorea producens*] | 40 | WP_070395084.1 |
| *atsH* | 415 | Glycosyltransferase | MULTISPECIES: glycosyl transferase family 1 [*Paenibacillus*] | 40 | WP_072329630.1 |
| *atsI* | 411 | Glycosyltransferase | glycosyl transferase family 1 [*Paenibacillus elgii*] | 41 | WP_108533365.1 |
| *atsJ* | 262 | *O-*Methyltransferase | hypothetical protein UV59_C0001G0033 [*Candidatus* Gottesmanbacteria bacterium GW2011_GWA1_43_11] | 41 | KKS86310.1 |
| *atsK* | 347 | Epimerase / Dehydratase | nucleoside-diphosphate sugar epimerase [*Candidatus* Staskawiczbacteria bacterium RIFOXYB1_FULL_37_44] | 59 | OGZ79612.1 |
| *atsL* | 287 | Nucleotidyltransferase | nucleotidyltransferase family protein [*Methermicoccus shengliensis*] | 47 | WP_042686739.1 |
| *atsM* | 405 | Glycosyltransferase | glycosyl transferase family 1 [*Alicyclobacillus acidiphilu*s] | 42 | WP_067621021.1 |
| *atsN* | 363 | Epimerase/ Dehydratase | NAD-dependent epimerase/dehydratase [*Chloroflexus aggregans* DSM 9485] | 72 | ACL24877.1 |
| *atsO* | 470 | MFS transporter | hypothetical protein CBC32_14015 [*Proteobacteria* bacterium TMED72] | 47 | OUU84711.1 |
| *atsP* | 371 | Regulator of protease activity | hypothetical protein C2W62_13455 [*Candidatus* 'Entotheonella serta'] | 64 | PON17388.1 |
| *atsQ* | 283 | Hypothetical protein | hypothetical protein [*Candidatus* 'Entotheonella palauensis'] | 32 | WP_034417458.1 |

**Table S7:** Putative vitamin B transporters as suggested by Rodionova et al.^3^ and their pairwise identity to ‘P. aureus’ CDSs.

| **Transporter** | **Predicted specificity,**  **B-vitamin pathway** | **Uniprot IDs** | **Pairwise identity [%]** |
| --- | --- | --- | --- |
| ThiX, ThiY, ThiZ | Thiamine, vitamin B_1_ | A9WDS0, A9WDR9, A9WDR8 | 44%, 24%, 42% |
| ThiW-EcfAA | Thiazole, vitamin B_1_ | A9WGB0, A9WGA9 | 25%, 0 |
| RibX, RibY | Riboflavin, vitamin B_2_ | A9WGD1, A9WGD2 | 46%, 38% |
| PanT, EcfT | Pantothenate, vitamin B_5_ | D6TJP8, D6TJP9, D6TJQ0 | 37%, 33% |
| BioY | Biotin, vitamin B_7_ | A9WAH3 | 0 |
| BtuF, BtuC, BtuD | Cobalamin, vitamin B_12_ | A9WCZ4, A9WCZ6, A9WCZ7 | 51%, 64%, 49% |

For

**Table S4:** Genome dataset for taxonomic and functional comparison.

**Table S6:** Vitamin B auxotrophy in ‘P. aureus’. Reactions with respective CDSs detected in the ‘P. aureus’ genome are highlighted in bold. All identifiers refer to entries in the KEGG database.

**Table S8**: Pfam-annotated CDSs in the ‘P. aureus’ TSY genome

, please refer to the supplementary document (Dataset S1).
